# HIV viral protein R induces loss of DCT1-type renal tubules

**DOI:** 10.1101/2023.02.02.526686

**Authors:** Khun Zaw Latt, Teruhiko Yoshida, Shashi Shrivastav, Amin Abedini, Jeff M. Reece, Zeguo Sun, Hewang Lee, Koji Okamoto, Pradeep Dagur, Jurgen Heymann, Yongmei Zhao, Joon-Yong Chung, Stephen Hewitt, Pedro A. Jose, Kyung Lee, John Cijiang He, Cheryl A. Winkler, Mark A. Knepper, Tomoshige Kino, Avi Z. Rosenberg, Katalin Susztak, Jeffrey B. Kopp

## Abstract

Hyponatremia and salt wasting is a common occurance in patients with HIV/AIDS, however, the understanding of its contributing factors is limited. HIV viral protein R (Vpr) contributes to HIV-associated nephropathy. To investigate the effects of Vpr on the expression level of the *Slc12a3* gene, encoding the Na-Cl cotransporter, which is responsible for sodium reabsorption in distal nephron segments, we performed single-nucleus RNA sequencing of kidney cortices from three wild-type (WT) and three Vpr-transgenic (Vpr Tg) mice. The results showed that the percentage of distal convoluted tubule (DCT) cells was significantly lower in Vpr Tg mice compared with WT mice (P < 0.05), and that in Vpr Tg mice, *Slc12a3* expression was not different in DCT cell cluster. The *Pvalb*^+^ DCT1 subcluster had fewer cells in Vpr Tg mice compared with WT (P < 0.01). Immunohistochemistry demonstrated fewer *Slc12a3*^+^ *Pvalb^+^* DCT1 segments in Vpr Tg mice. Differential gene expression analysis comparing Vpr Tg and WT in the DCT cluster showed *Ier3*, an inhibitor of apoptosis, to be the most downregulated gene. These observations demonstrate that the salt-wasting effect of Vpr in Vpr Tg mice is mediated by loss of *Slc12a3*^+^ *Pvalb*^+^ DCT1 segments via apoptosis dysregulation.

## Introduction

The distal convoluted tubule (DCT) is the shortest segment of the distal nephron. Its physiological roles include maintaining sodium homeostasis. The DCT specifically expresses the thiazide-sensitive NCC (sodium-chloride cotransporter), encoded by *SLC12A3,* and is responsible for the reabsorption of luminal Na^+^ into DCT cells.

The DCT shows vulnerability as well as adaptability to conditions that interfere with Na reabsorption. In a study by Kaissling and colleagues, minipumps containing furosemide were implanted in rats for six days. The rats drank large volumes of salt solution provided *ad libitum* to compensate for urine sodium losses. This intervention resulted in the enlargement of kidney cortices, with histologic evidence of proliferation of distal convoluted tubules (1).

In 1996, Loffing and colleagues showed that in rats given thiazide, the DCT epithelium lost the structural characteristics of electrolyte-transporting epithelia, with cells in various stages of apoptosis. Thiazide-sensitive sodium-chloride cotransporter (rTSC1) trptanscripts were greatly reduced in kidney cortex homogenates and were almost entirely absent in damaged DCT cells. Focal inflammatory cell infiltrates were localized to the DCT, whereas all other tubular segments were unaffected by this intervention. Thus, inhibition of apical NaCl entry into DCT cells by thiazides is associated with apoptosis of DCT cells and associated inflammation (2).

HIV-associated nephropathy (HIVAN) manifests dysfunction and pathology of both glomeruli and tubules. In the past, attention has been particularly focused on podocytopathic injury of HIVAN glomerulopathy. Recently, HIV-associated tubular injury has drawn increased attention, as hyponatremia due to sodium wasting (3) is common in hospitalized HIV/AIDS patients (4, 5).

Previously, we demonstrated a reduction of NCC protein in the cortex of HIV viral protein R (Vpr) transgenic mice. These mice showed increased urine sodium loss, despite being on a low sodium diet for 4 days. The reduction in NCC protein levels in Vpr Tg mice was mediated by reduced expression of its encoding gene (*Slc12a3*) in DCT cells (6).

It is known that HIV-1 Vpr causes renal cell dysfunction and contributes to HIVAN pathogenesis (3) as well as DCT dysfunction (6). We wished to explore mechanisms whereby Vpr compromises DCT function by single-nucleus RNA sequencing (snRNA-seq) on kidney cortices of HIV-transgenic mice maintained on a sodium-deficient diet. Our goal was to characterize, at the single-cell resolution, the molecular and the cell biological effects of Vpr across the nephron, particularly, on distal tubular cells.

## Results

### DCT cell number was reduced in Vpr Tg mice

We performed snRNA-seq analysis of three WT and three Vpr Tg mouse kidney cortex samples. Unsupervised clustering in Seurat gave 25 cell clusters which were annotated using known canonical cell type marker genes: eight proximal tubule cell clusters (labeled as PT-S1,PT-S1_UK, PT-S1/S2, PT-S2, PT-S2/S3, PT-S3, Glyco and Mito), three distal tubular cell clusters (DCT expressing *Slc12a3*, CNT expressing *Slc8a1* and PC expressing *Aqp2*), two endothelial cell clusters (cortical EC and vascular EC), two intercalated cell clusters (IC-A and IC-B), two thick ascending limb clusters (TAL-1 and TAL-2), and one cluster each of podocytes, mesangial cells (MC), smooth muscle cells (SMC), macula densa cells, thin descending limb (tDL), fibroblasts, T cells and other immune cells (**Figure 1**).

**Figure 1.**
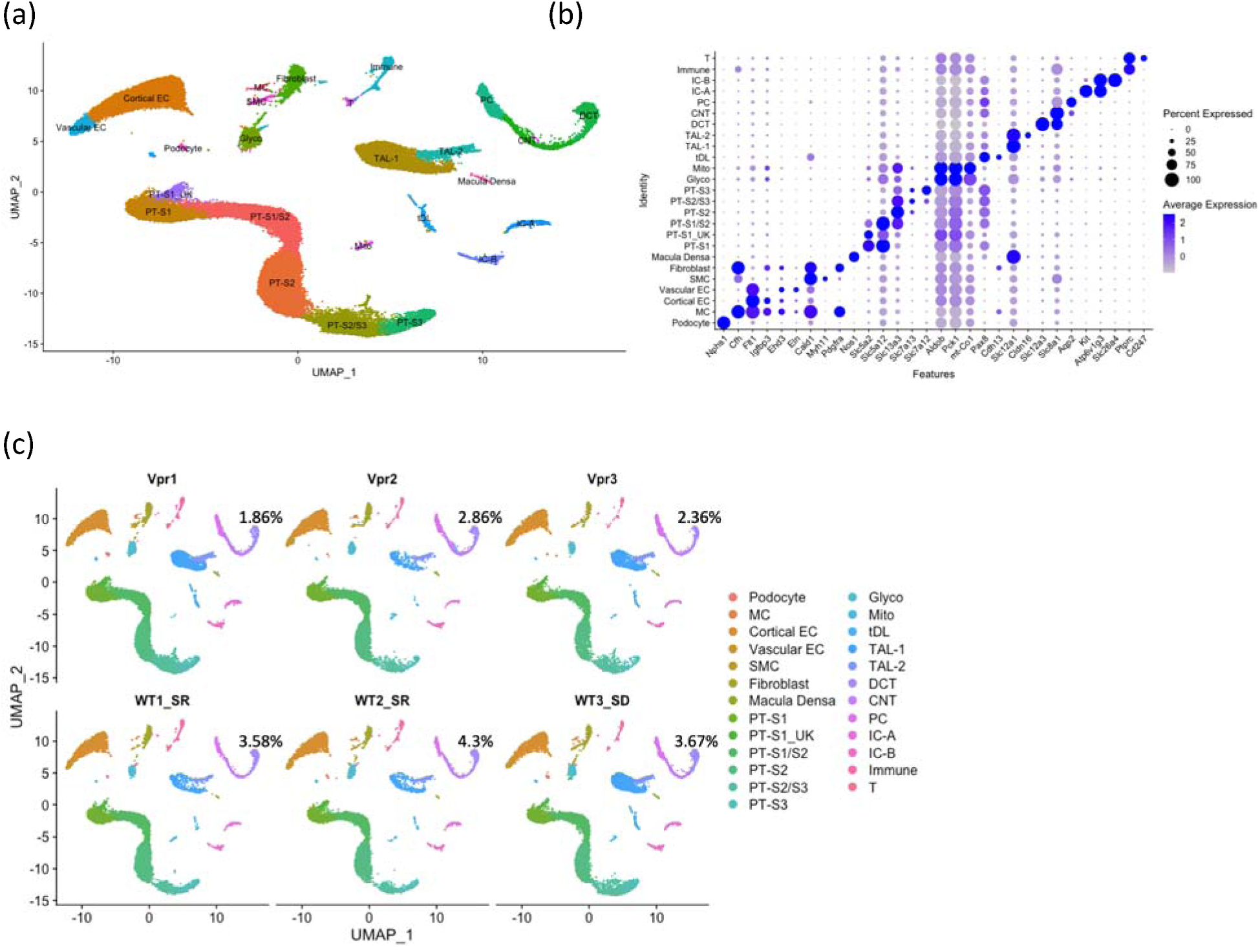
snRNA-seq of mouse kidney cortex samples. (a) UMAP projection of 10x datasets from six samples after anchor-based integration. (b) Dot plot showing the marker genes to identify the cell type in each cluster: podocyte; mesangial cell (MC); cortical endothelial cell (Cortical EC); vascular endothelial cell (Vascular EC); smooth muscle cell (SMC) ; fibroblast; macula densa; proximal tubule, S1 segment (PT-S1); proximal tubule, S1 segment of unknown subtype (PT-S1_UK); proximal tubule, S1 and S2 segments (PT-S1/S2) ; proximal tubule, S2 segment (PT-S2); proximal tubule, S2 and S3 segments (PT-S2/S3); proximal tubule, S3 segment (PT-S3); Cells enriched in glycolytic enzymes (Glyco); Cells enriched in mitochondrial transcripts (Mito); thin descending limb (tDL); thick ascending limb 1 (TAL-1); thick ascending limb 2 (TAL-2) ; distal convoluted tubule (DCT) ; connecting tubule (CNT); principal cell (PC); type A intercalated cell(IC-A); type B intercalated cell (IC-B); immune cell (Immune) ; T cell (T). (c) UMAP projections of individual sample showing the percentages of DCT cells to overall cells in each sample.

For each sample, we calculated the percentages of each cell type as a function of total cell number. The WT and Vpr Tg samples showed similar cell percentages across most of the identified renal epithelial cell clusters. A notable exception was found in the DCT cluster, which showed a reduced cell fraction in the Vpr Tg samples (1.86 %, 2.86% and 2.36% (2.36±0.5%) in each of the Vpr Tg samples as compared with 3.58 % and 4.3% and 3.67% (3.85±0.4%) in WT samples, respectively) (P < 0.05) (**Figure 1c** **and Supplementary Table 1**).

### The CNT cluster showed the highest expression of *Nr3c2* and *Hsd11b2*, genes essential for aldosterone responsiveness

Vpr has been reported to bind the mineralocorticoid receptor (MR, encoded by *Nr3c2*) and to inhibit MR-mediated upregulation of *Slc12a3* by acting as MR corepressor (6). The *Hsd11b2* gene encoding the enzyme 11-β-hydroxysteroid dehydrogenase 2, is required to metabolize glucocorticoids which are present in higher concentrations within cells, compared to aldosterone. High expression levels of *Hsd11b2* promote specific binding of mineralocorticoids to the MR rather than the glucocorticoid receptor (GR).

To investigate the cell types most responsive to aldosterone, we examined the expression profiles of *Nr3c2* and *Hsd11b2* in distal tubular cell clusters. Expression levels of *Nr3c2* and *Hsd11b2* were highest in CNT and PC clusters, while their expression levels were very low in the DCT cluster (**Figure 2a**). There was little overlap in the expression profiles of *Slc12a3* and of these two genes (**Figure 2b** and **2c**). The differential expression test also showed no downregulation of *Slc12a3* gene in the DCT cluster (**Figure 2d**).

**Figure 2.**
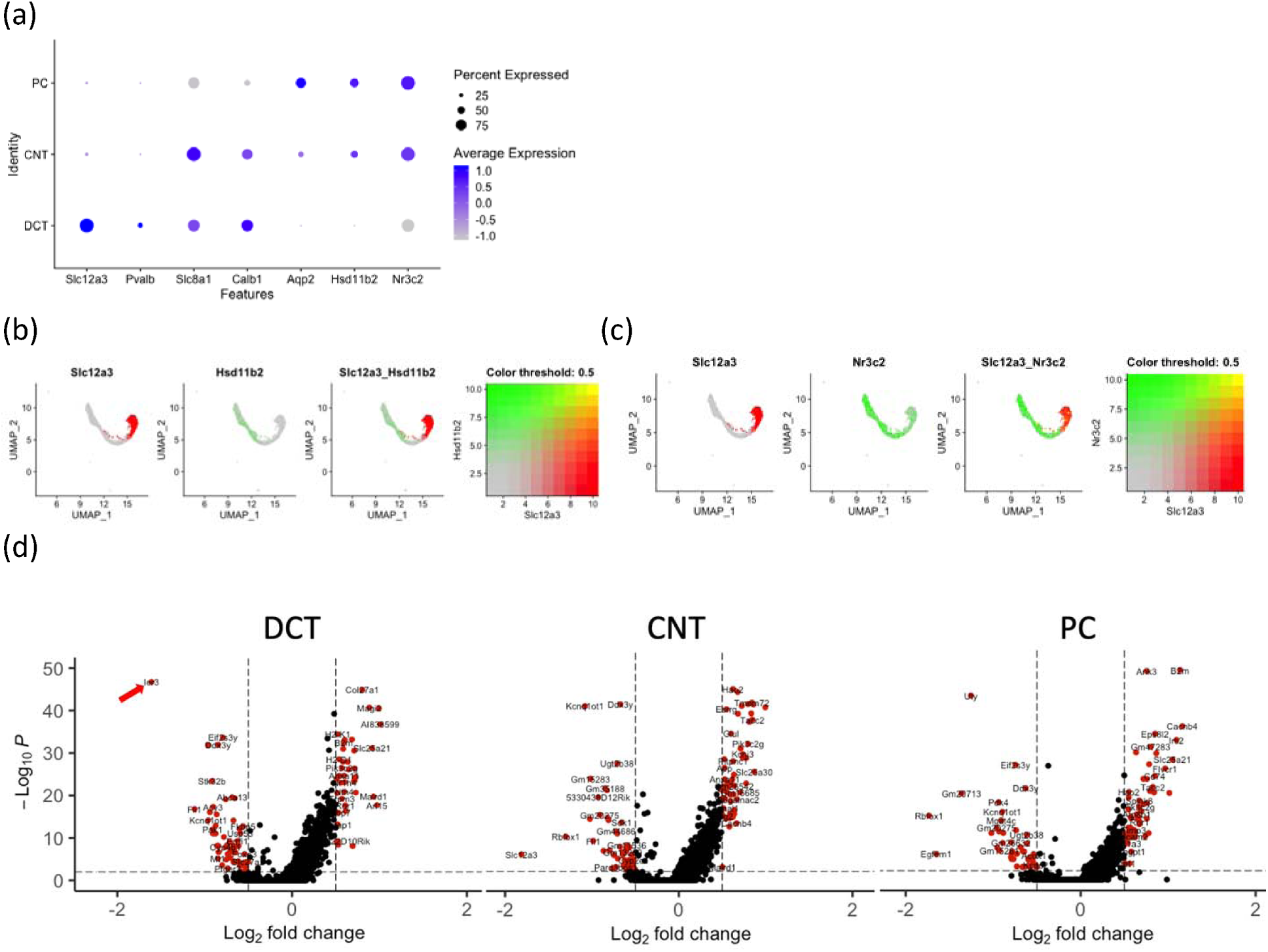
Expression pattern of *Slc12a3*, *Nr3c2*, *Hsd11b2* and other genes across distal tubular clusters. (a) Dot plot showing the expression of genes distinguishing distal tubular cell types. (b) Feature plots of individual samples showing the expression overlap between *Slc12a3* and *Hsd11b2*. (c) Feature plots showing the overlap of expression between *Slc12a3* and *Nr3c2*. (d) Volcano plots showing differentially expressed genes in DCT, CNT and PC clusters. Red arrow indicates *Ier3* downregulation in DCT cluster.

### A DCT-specific open chromatin mark spans the *Slc12a3* region

*Slc12a3* is a DCT-defining signature gene and it is known from single-nucleus assay for accessible chromatin-sequencing (ATAC)-seq data that there is an open chromatin over this gene region in DCT cells. Using human and developing mouse kidney ATAC-seq data from the Susztaklab Kidney Biobank (https://susztaklab.com/) (7–15), we observed that the open chromatin feature spanning over 50 kb is present in the *SLC12A3*/*Slc12a3* region in a DCT-specific manner in both human and mouse genomes. This suggests that this region may qualify as a super-enhancer that ensures robust expression of lineage-defining *Slc12a3* in DCT cells (**Figure 3**).

**Figure 3.**
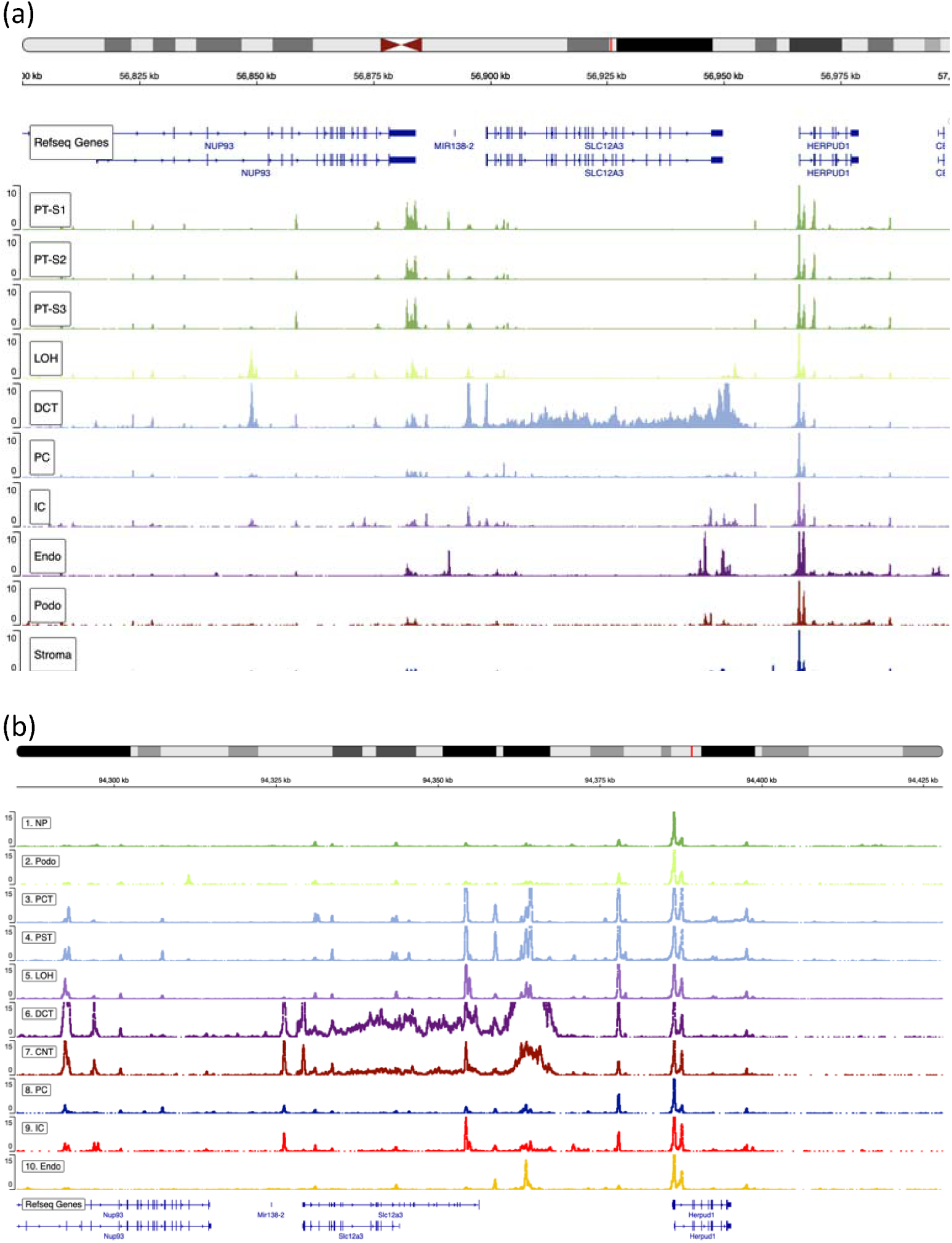
snATAC-seq data of kidney cells showing transposase-accessible open chromatin peaks around *Slc12a3* regions retrieved from Susztaklab Kidney Biobank. (a) Open chromatin regions around *SLC12A3* region on chromosome 16 of the human kidney. (b) Open chromatin regions around *Slc12a3* region on chromosome 8 of the developing mouse kidney.

To look for the evidence of the regulatory association between the open chromatin region and *SLC12A3*/*Slc12a3* gene expression, we inspected the expression quantitative trait loci (eQTL) and the H3K27ac epigentic histone acetylation mark in human *SLC12A3* region from the Genotype Tissue Expression (GTEx) portal and candidate cis-regulatory elements (cCRE) data from the Encyclopedia of DNA Elements (ENCODE) databases. The GTEx data showed multiple eQTL SNPs associated with the *SLC12A3* expression levels across the open chromatin data (**Supplementary Table 2**) and H3K27ac enhancer mark in frontal cortex, left ventricle, skeletal muscle and lung (**Supplementary Figure 3**). In the ENCODE database, we sought evidence of regulatory elements in this region, with a focus on glomerular epithelial cells and kidney tissues from human subjects (**Supplementary Figure 4**). There were multiple cCREs across the region, identified in all registry samples. Furthermore, there were CTCF ChIP-seq signals for the CCCTC-binding factor (CTCF), a highly conserved zinc finger protein and transcription factor, present in the kidney tissue data from three male adult subjects (**Supplementary Figure 4b**).

### Vpr Tg mice showed loss of *Pvalb^+^* DCT1 cell fraction

We performed sub-clustering of the three distal tubule cell clusters (DCT, CNT and PC) to pinpoint the cell sub-populations that were lost in the Vpr Tg mouse and to identify the underlying pathways involved in the cell loss. This approach identified seven subclusters (three DCT clusters and two clusters each for CNT and PC) (**Figure 4**). Of the three DCT clusters in Vpr Tg samples, a decreased cell fraction was observed only in the *Pvalb*^+^ DCT1 cluster (Cluster 0) (1%, 1.93% and 1.47% in Vpr Tg and 3.03%, 3.41%, 2.87% in WT samples, P < 0.01) but not in *Slc8a1*^+^ *Calb1*^+^ DCT2 cluster (Cluster 5) or in *Nr3c1*^+^ DCT cluster (Cluster 6) (**Supplementary Table 1**). The DCT subclusters we identified are consistent with those described in a recent report employing targeted single-cell RNA-seq on distal nephron cells (16).

**Figure 4.**
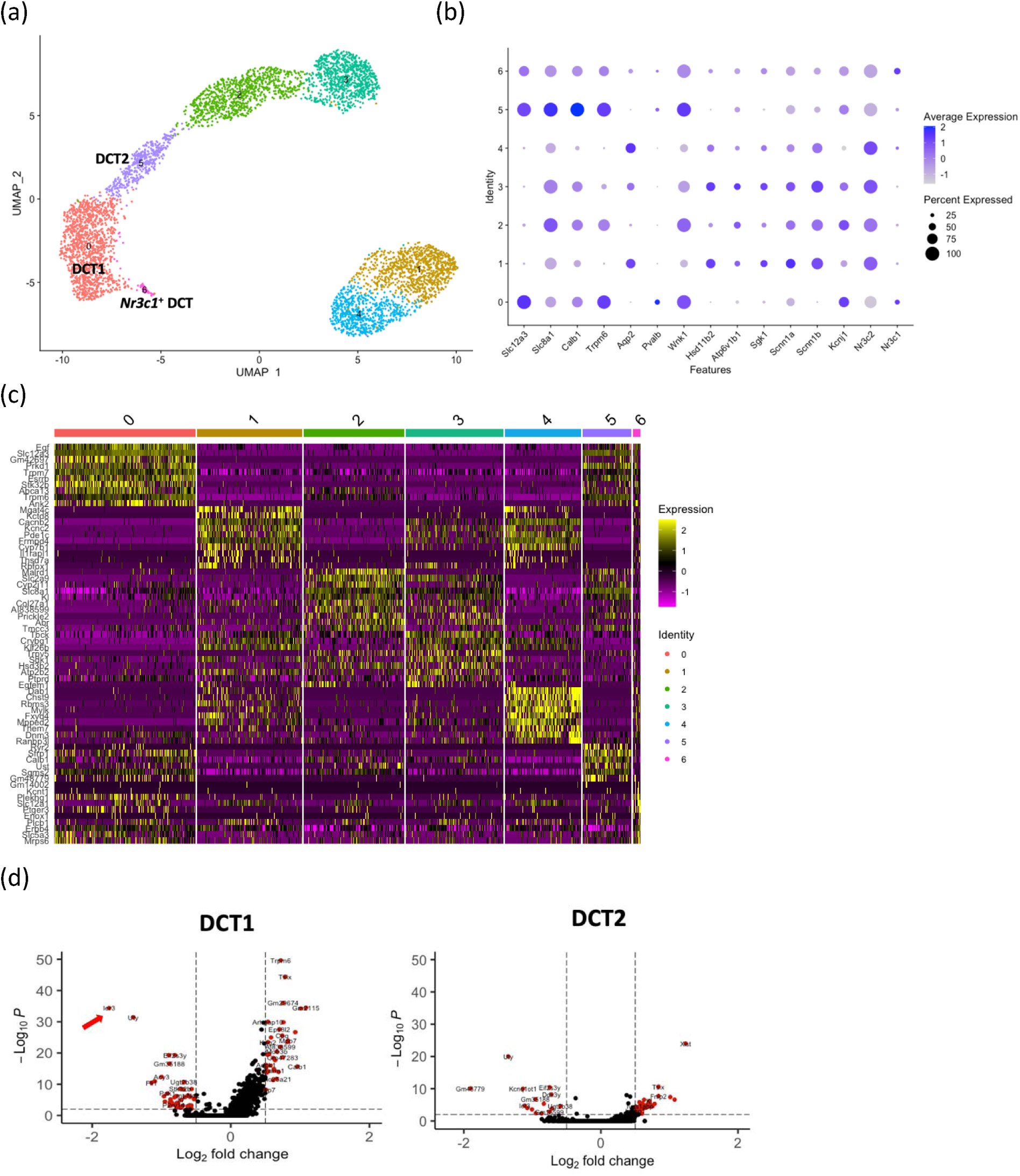
Subclustering of distal tubule clusters (DCT, CNT and PC). (a) UMAP plot showing 7 subclusters. (b) Dot plot showing marker genes to identify sub-populations of distal tubular cells. (c) Heatmap showing the ten highly expressed genes in each subcluster compared to all remaining subclusters. (d) Volcano plots showing downregulation of *Ier3* gene in DCT1 but not in DCT2 subcluster.

### Downregulation of anti-apoptotic *Ier3* gene in DCT1 cells of Vpr Tg mice

Differential expression analyses were performed in each distal tubular cell subpopulation. Differential expression tests showed downregulation of anti-apoptotic *Ier3* gene, specifically in DCT1 but not in DCT2 subcluster (**Figure 4d**). Therefore, the *Ier3* downregulation observed in DCT cluster (**Figure 2d**) was driven by the DCT1 subpopulation. Other distal tubular cell (CNT and PC) subclusters did not show downregulation of *Ier3*.

### Immunostaining studies confirmed the DCT1 cell loss and the increased apoptotic rates in Vpr Tg mice

To confirm the results obtained in the snRNA-seq analysis, *in situ* hybridization (ISH), immunohistochemistry (IHC) and terminal deoxynucleotidyl transferase-mediated dUTP nick end labeling (TUNEL) apoptosis assay were performed.The results were compared between WT and Vpr Tg mice. ISH showed marked reduction of *Slc12a3* and *Pvalb* transcripts in Vpr Tg mouse cortex (**Figures 5a-c** **and Supplementary Figure 5**), suggesting the loss of DCT1 cells. The IHC staining and quantification of DCT1-specific parvalbumin showed a significant decrease in the percent of parvalbumin^+^ cortical tubular area in Vpr Tg samples (**Figures 5d and 5e**). Vpr Tg samples had more TUNEL-positive nuclei compared with WT samples (**Figures 5f and 5g**) although TUNEL-positivity was not specific to DCT1 cells. Thus, it is likely that reduced *Slc12a3* and *Pvalb* expression observed in Vpr Tg mouse cortex is caused by induction of apoptosis and subsequent loss of DCT1 cells.

**Figure 5.**
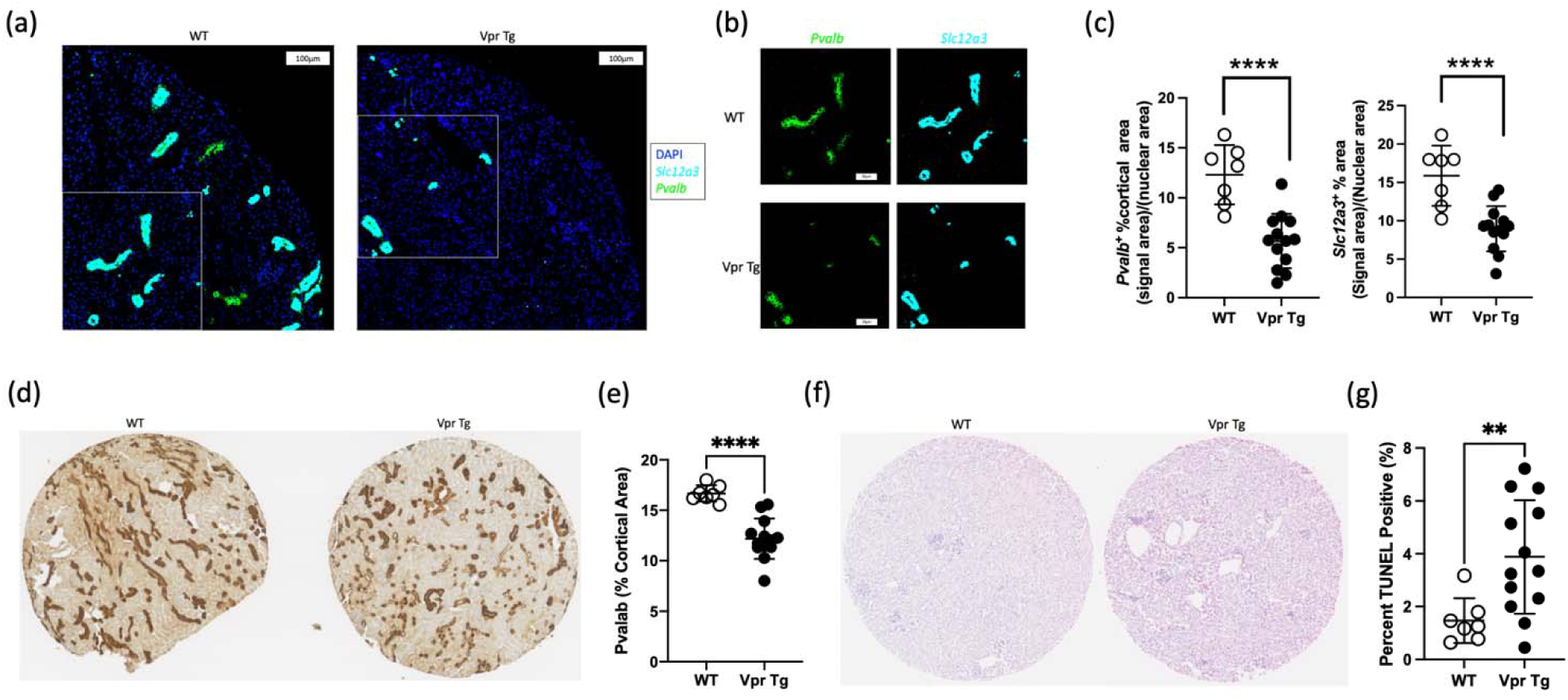
Imaging results of WT and Vpr Tg kidney cortex samples. (a) RNAScope images of a quarter of a tissue microarray section showing that Vpr mice had less *Slc12a3* and *Pvalb* fluorescence signals than WT mice. (b) Areas of images from Figure 5a marked by the white rectangles showing individual genes for *Pvalb* and *Slc12a3*. (c) Dot plots showing unpaired t-tests comparing the cortical areas positive for fluorescence signals of *Pvalb* and *Slc12a3* between WT and Vpr Tg mice. (d) Tissue microarray sections showing reduced parvalbumin (Pvalb) staining in Vpr Tg mice compared to WT (e) Dot plot of the unpaired t-test comparing parvalbumin (+) cortical area between seven WT mouse and thirteen Vpr Tg mouse samples. (f) Tissue microarray sections showing TUNEL staining of WT and Vpr Tg samples. (g) Dot plot showing the results of MannWhitney U test comparing the percent of TUNEL (+) nuclei between WT and Vpr Tg samples. The Pvalb and TUNEL values of each sample on Y-axis were average values from triplicates.

### DCT cell loss *in vitro*

Next, to confirm the DCT cell loss due to Vpr, we used the 209/Mouse DCT (mDCT) cells (originally immortalized using adenovirus). These cells were exposed to soluble Vpr for twenty-four hours and then processed for the flow cytometric analysis. The results showed a significant increase in propridium iodide-positive dead cells as compared to cells without exposure to Vpr (**Figures 6a and 6b**). The qPCR results of Vpr-exposed mDCT cells also showed a significant downregulation of anti-apoptotic *Ier3* gene (**Figure 6c**) consistent with the in vivo Ier3 downregualtion in Vpr Tg mice (**Figures 2d and 4d**).

**Figure 6.**
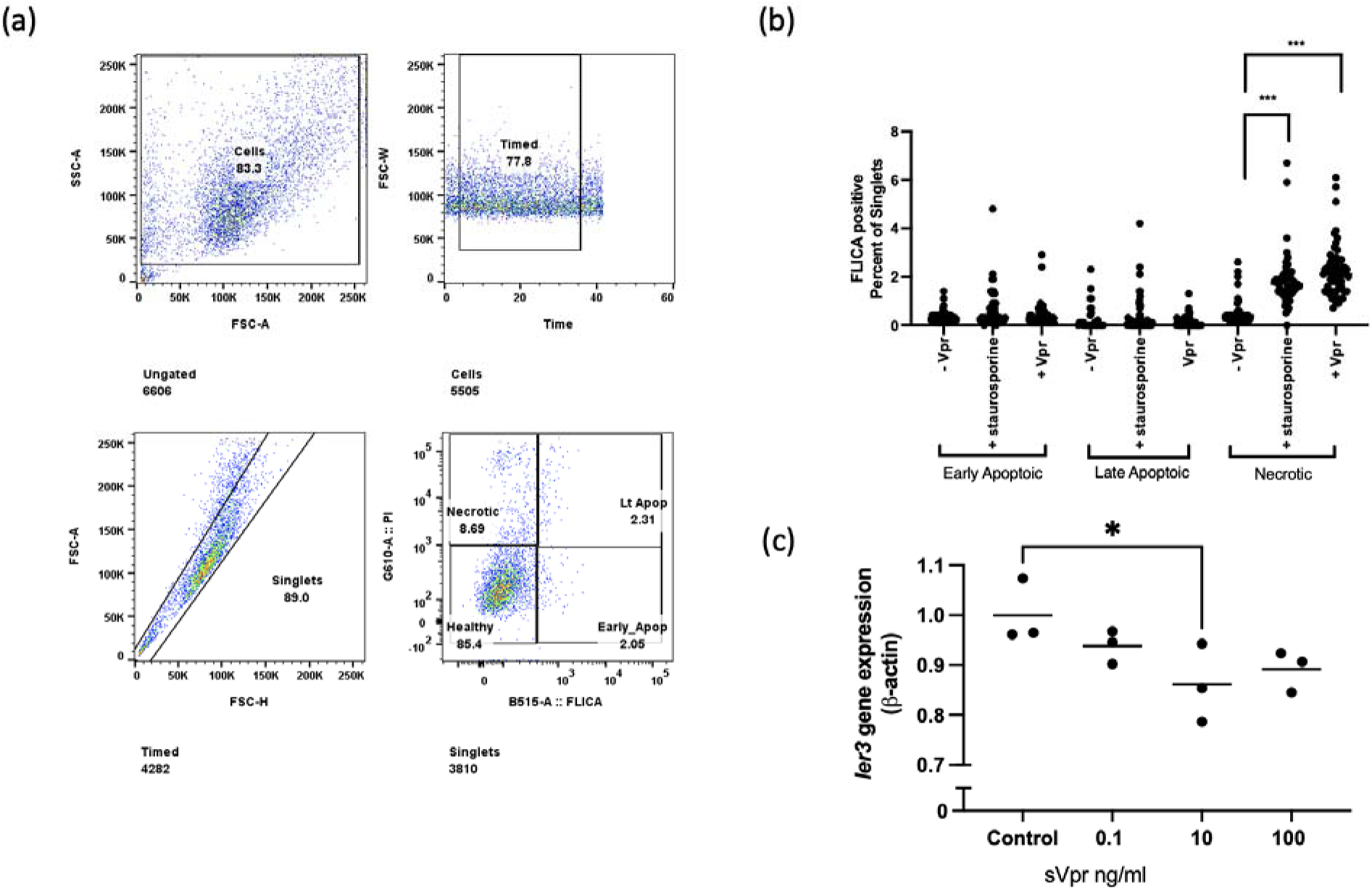
*In vitro* detection of apoptosis and *Ier3* downregulation in sVpr treated 209 mDCT cells. (a) Flow cytometry of apoptotic markers (FLICA and PI) on mDCT cells treated with Vpr. Propidium iodide (PI) was used as the live/death discriminator. (b) Increase (p < 0.001) in PI-labelled nuclei representing apopotic mDCTcells after treatment with either 1ng per well of sVpr or 0.1uM of positive control staurosporine for 24 hours, compared to untreated controls. (c) The qPCR analysis showed significant downregulation of *Ier3* mRNA at 10 ng/ml of Vpr, compared to untreated controls.

### DCT cell loss in Pax8-Vpr mouse model

To verify the DCT1 cell loss caused by Vpr in an independent model, we analysed the published Pax8-Vpr mouse snRNA-seq dataset generated by Chen et al. (17). Distal tubular cell clusters expressing *Slc12a3* (DCT), *Slc8a1*(CNT) and *Aqp2* (PC) were selected for subclustering analysis and cellular percentages were compared between WT and Pax8-vpr mice. We found that the *Slc12a3*^+^ *Pvalb*^+^ DCT1 cell percentages were lower in Pax8-vpr mice compared to WT mice (**Figure 7**), similar to the observation of DCT1 cell loss in Pepck-Vpr mice.

**Figure 7.**
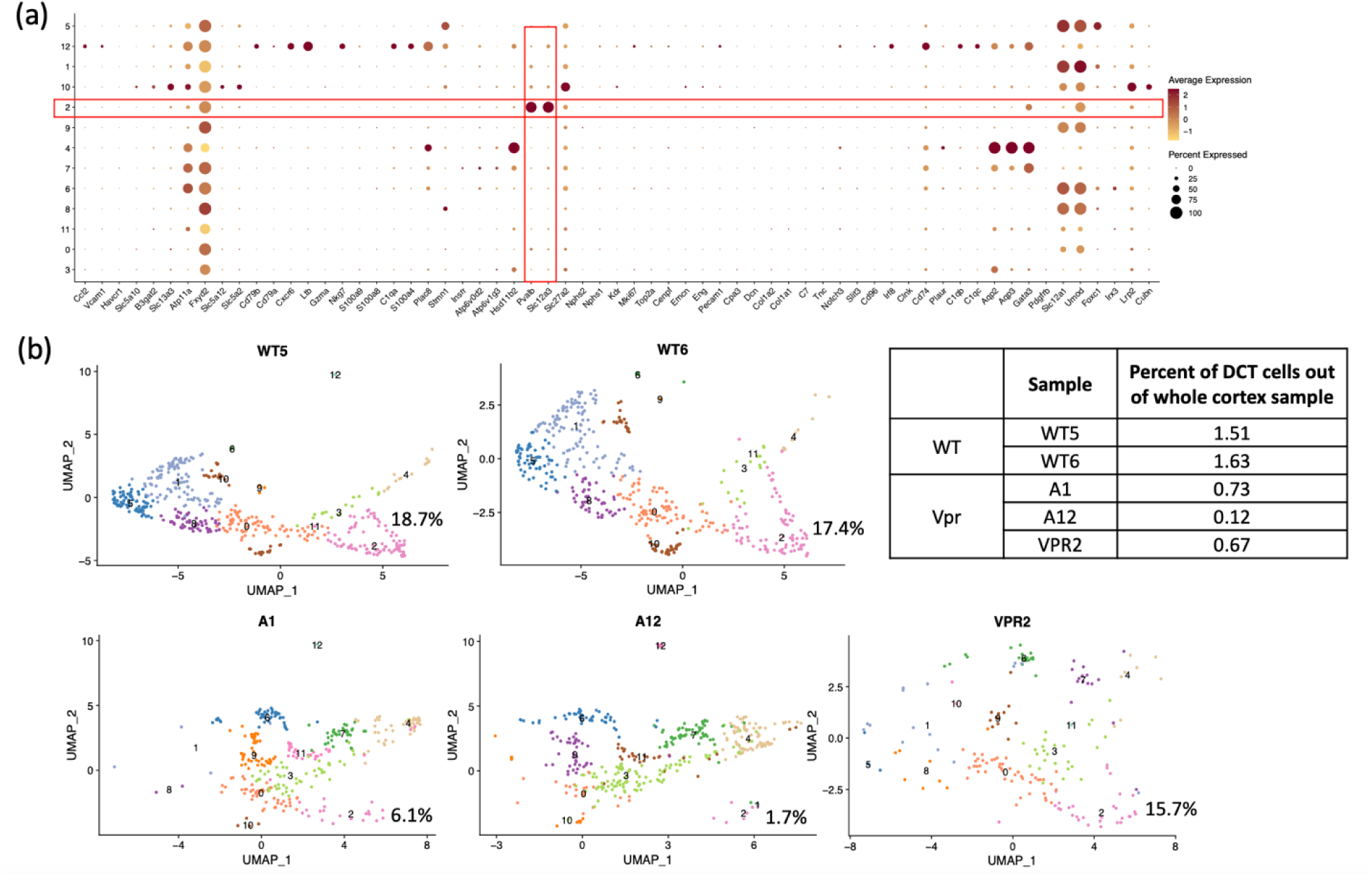
Loss of DCT1 cells in Pax8-vpr mice. (a) Subclustering of distal tubular cells identified *Slc12a3*^+^ *Pvalb*^+^ DCT1 subcluster 2. (b) UMAP plots showing percentages of subcluster 2 (DCT1 cells) out of all distal tubular cells, and table showing percentages of DCT1 cells out of the whole cortex samples.

## Discussion

Various interventions in mice, including thiazide administration (2), deletion of *Slc12a3* or *Pvalb* (18, 19), over-expression of WNK4 (20), or loss of Ste20-related proline-alanine-rich kinase (SPAK) (21), have been shown to reduce the levels of NCC protein and induce shortening of DCT length.

We recently reported that transgenic Vpr expression causes downregulation of NCC protein expression in renal distal tubules and proposed that binding of Vpr to the mineralocorticoid receptor (MR) blocks aldosterone/MR-mediated transcription of *Slc12a3* in mice (6). Therefore, to investigate the molecular mechanisms underlying the reduction of NCC expression, we applied single-nucleus RNA sequencing to WT and Vpr Tg mouse kidney cortices.

Compared with WT mice, Vpr Tg mice showed reduced fractions of *Pvalb*^+^ DCT1 cells. The most significantly downregulated gene in DCT1 cells was *Ier3*, which regulates apoptosis in a variety of cells and organs and has been reported to have anti-apoptotic effect in cardiomyopathy and some cancers (22–29). The in vitro treatment of mouse DCT (mDCT) cells with soluble Vpr also showed similar downregulation of *Ier3*. Vpr is known to induce apoptosis through the cell-cycle arresting effect (3). However, when we evaluated the aggregate scores of cell cycle phase-specific genes in WT and Vpr Tg samples, the scores of G2M- and S phase-specific genes were similar between the WT and Vpr Tg mice (**Supplementary Tables 3 and 4** and **Supplementary Figure 6**) suggesting that, at least in the present setting, the cell cycle arrest function of Vpr may not contribute to its transcriptional effects. Similar DCT1 cell loss in Pax8-vpr mouse model lends support to the data presented here.

Interestingly, there was no significant downregulation of *Slc12a3* gene expression in DCT cells. Aldosterone upregulates the expression of NCC through augmenting the transcriptional activity of the MR on the *Slc12a3 gene*. This aldosterone action is also dependent on the intracellular expression of the enzyme *Hsd11b2,* which metabolizes glucocorticoids (specifically corticosterone in mice) into inactive 11-dehydroxycortisone. Glucocorticoids have higher intracellular concentrations than aldosterone and compete with aldosterone for binding to MR (30, 31). In the present study, the highest expression of MR (*Nr3c2*) and *Hsd11b2* was found in the CNT. The expression levels of these genes in DCT cell cluster is relatively low. These findings are consistent with those of Chen *et al.,* who also observed higher expression of these genes in DCT2 and CNT than in other nephron segments (16). This pattern of gene expression suggests that DCT1 is not a major site of aldosterone action and Vpr inhibition of MR most likely does not occur in mouse DCT1 cells.

Using Susztaklab Kidney Biobank data, we found that the *Slc12a3* region spanning over 50 kb is characterized as open chromatin, specifically in DCT. The GTEx and ENCODE data showing the cis-regulatory elements, especially the CCCTC-binding factor (CTCF)-binding sites in adult kidney tissue, further suggests that this region may act as a super-enhancer. The CTCF was reported to be bound near and within some super-enhancers (32–35), and is critical for super-enhancer-mediated gene expression regulation (36–38).

In the DCT, *Slc12a3* is the cell type-defining gene and therefore, the open chromatin region around this gene may function as a super-enhancer. Super-enhancers are usually larger than other enhancer regions which facilitate the binding of multiple key transcription factors and promote cellular development/differentiation into particular cell lineages (39–41). Taken together, these findings suggest that DCT cells have a strong basal expression of *Slc12a3,* which is less dependent on aldosterone and more resistant to inhibitory factors such as Vpr.

In conclusion, we applied the snRNA-seq approach to analyze kidney cortices from Vpr Tg mice, a model of salt wasting in HIV-associated kidney disease, to gain further insight into the disease mechanisms. The results indicate that the HIV-1 Vpr induced DCT1 apoptosis and downregulated *Ier3*. In addition, we detected similar DCT1 cell loss in Pax8-Vpr mouse model, landing substantial support to the role of Vpr in DCT1 cell loss. Taken together, these findings have extended our understanding of salt wasting mechanisms in HIV/AIDS. Further studies will be required to clarify mechanisms underlying the *Ier3* downregulation and apoptosis in DCT1 cells.

## Methods

### Generation and maintenance of Vpr transgenic mice

Detailed information about the generation, characterization, and genotyping of tetracycline repressible Vpr transgenic mice have been described previously (6, 42), and in the Supplementary materials.

### Preparation of mouse kidney cortex samples for snRNA-seq

We selected cortex samples from three Vpr Tg mice and one WT mouse that were fed with low sodium diet for four days (6). We also performed snRNA-seq on two WT kidney cortex samples from FVB mice maintained on regular laboratory chow. Nuclei from frozen mouse kidney outer cortex tissue samples were prepared following the protocol of Kirita *et al*. (43). Briefly, ∼ 8 mm^3^ tissue fragments were cut by razor blade in EZlysis buffer (#NUC101-1KT, Sigma, Darmstadt, Germany) and first homogenized 30 times with a Dounce homogenizer equipped with a loose pestle and homogenized 5 times with a tight pestle. After 5 min of incubation, the homogenate was passed through a 40 µm filter (PluriSelect, El Cajon, CA) and centrifuged at 500xg at 4°C for 5 min. The pellet was washed with EZlysis buffer (Sigma Aldrich, St Louis, MO) and again centrifuged at 500x g at 4°C for 5 min. The pellet was resuspended in Dulbecco’s phosphate-buffered saline supplemented with 1% fetal bovine serum and passed through a 5 µm filter (PuriSelect, El Cajon, CA) for obtaining the final nuclear suspension.

The prepared nuclei were loaded onto a 10X Chromium Chip G (10X Genomics, San Francisco, CA) for formation of gel beads in emulsion (GEM). Single nuclear isolation, RNA capture, cDNA preparation, and library preparation were accomplished following the manufacturer’s protocol (Chromium Next GEM Single Cell 3’ Reagent Kit, v3.1 chemistry, 10x Genomics, Pleasanton, CA).

The prepared cDNA libraries were sequenced at the Frederick National Laboratory for Cancer Research Sequencing Facility (NCI, Frederick, MD). The mean reads per cell for WT1, WT2, WT3, Vpr1, Vpr2 and Vpr3 samples were 28,731, 51,350, 75,706, 48,460, 93,558 and 76,878, respectively, and the Q30 bases in the RNA were 90.9%, 93.4%, 92.5%, 93.9%, 93.1% and 93%, respectively. Analysis was performed with the Cell Ranger v5.0.0 software using the default parameters including the pre-mRNA analysis (10x Genomics, Pleasanton, CA). The reference was built from the mm10 (*Mus musculus*) reference genome complemented with reported HIV-1 viral sequences.

### snRNA-seq data analysis

The Cell Ranger output files of individual 10x Genomic datasets were used as input for SoupX (44) (version 1.6.1) (https://github.com/constantAmateur/SoupX) to estimate and correct the contamination fraction for ambient or cell-free RNA released during the process of lysing cells for nuclear preparation. The 10x datasets corrected for ambient RNA were converted into Seurat objects by excluding nuclei with (a) less than 200 or more than 4000 genes, (b) total RNA count of more than 15000, and (c) more than 20% of mitochondrial transcripts (Seurat version 4.1.1) (45). The data were adjusted using the mitochondrial transcript percentage of each nucleus, based on the 2000 most variable nuclear-encoded genes.

Potential cell doublets were predicted and removed by DoubletFinder (46) (version 2.0.3) (https://github.com/chris-mcginnis-ucsf/DoubletFinder) by assuming a doublet rate of 7.6% and using ten principal components, pN and pK values of 0.25 and 0.005, respectively (**Supplementary Figure 1**). The three datasets (WT, **WT_SR** and Vpr Tg) were integrated using an anchor-based integration. After batch correction using Harmony (47) (version 0.1.0) (https://github.com/immunogenomics/harmony) (**Supplementary Figure 2**), clustering was done with 30 principal components and a resolution of 0.5. Subclustering of distal tubule data was performed by subsetting three distal tubule cell clusters – DCT, CNT and principal cell (PC) through identifying the most variable genes in these three clusters in each sample, followed by anchor-based integration, and by repeating batch correction. The differential expression tests between cells from WT and Vpr Tg samples were performed using Wilcoxon rank sum tests implemented in Seurat package.

Detailed methods of tissue microarray preparation, *in situ* hybridization, confocal microscopy and image analysis, immunohistochemistry (IHC) of parvalbumin, TUNEL assay to detect apoptotic cells and quantification of IHC and TUNEL images and caspase assay and quantitative PCR of *Ier3* using 209 mDCT cells are described in supplementary materials.

## Abbreviations used

DCT: distal convoluted tubule
Vpr: viral protein R
HIVAN: HIV-associated nephropathy
snRNA-seq: single nucleus RNA sequencing

## Acknowledgment

We thank the Sequencing Facility and Bioinformatics Group (Frederick National Laboratory for Cancer Research (FNLCR), NCI, NIH) for sequencing and informatics support. This work utilized the computational resources of the NIH HPC Biowulf cluster (http://hpc.nih.gov). The project has been supported in part by the National Institutes of Health and the National Cancer Institute Intramural Research Program (CAW) and under contract 75N91019D00024. The authors also thank the NIDDK Advanced Light Microscopy & Image Analysis Core (ALMIAC) and the NHLBI FACS core. We acknowledge Huiyan Lu, Kris Ylaya for technical support, and Luis Fernando Menezes and Noor Khalil for critical manuscript review.

The content of this publication does not necessarily reflect the views or policies of the Department of Health and Human Services, nor does mention of trade names, commercial products, or organizations imply endorsement by the U.S. Government.

**The GEO accession numbers and tokens of the datasets used in this study were shared with the editors.**

## Author contribution

KZL, SS and JBK conceived and designed the study. SS and TY performed mouse experiments. TY, SS and KZL did the snRNA-seq experiments. KZL, TY, SS, AZR and JBK did analysis and interpreted results. AA and KS shared ATAC-seq data and gave critical comments. YZ helped with initial analysis of sequencing data. JYC, SH and JMR helped with ISH imaging experiments and quantification analysis. SS and PD helped with FACS analysis and interpretation of mDCT cells. ZS, KL and CH did the analysis of Pax8-vpr mice. MAK, HL, PAJ, JH, CAW and TK gave critical comments and suggestions. KZL and SS wrote the manuscript.

## Supplementary Materials

## Supplementary Methods

## Supplementary Figures

**Supplementary Figure 1.** Results of doublet detection analysis in each sample by DoubletFinder.

**Supplementary Figure 2.** SnRNA-seq analysis of the dataset after anchor-based integration and batch correction by harmony.

**Supplementary Figure 3.** H3K27ac enhancer mark on human chromosome 16 covering *SLC12A3* region in frontal cortex, left ventricle, skeletal muscle and lung retrieved from GTEx database.

**Supplementary Figure 4.** Candidate cis-regulatory elements (cCREs) in human *SLC12A3* region retrieved from ENCODE database showing multiple enhancer-like elements (yellow vertical bars) in all registry samples.

**Supplementary Figure 5.** Imaging results of WT and Vpr Tg and cortex samples.

**Supplementary Figure** 6. Violin plots showing the aggregate gene scores of cell cycle genes in distal tubular cell subclusters.

## Supplementary Tables

**Supplementary Table 1**. Metrics of the 25 cells clusters and 3 DCT sub-clusters that were identified in the unsupervised clustering of six mouse renal cortex samples.

**Supplementary Table 2**. Expression quantitative trait locus (eQTL) SNPs associated with *SLC12A3* expression levels across different tissues retrieved from the Genotype Tissue Expression (GTEx) database.

**Supplementary Table 3.** The mouse G2M phase genes used in the aggregate gene scores shown in **Supplementary Figure 5a**.

**Supplementary Table 4.** The mouse S phase genes used in the aggregate gene scores shown in Supplementary Figure 5b.

