## Supplementary Materials for "HIV viral protein R induces loss of DCT1-type renal tubules"

### Supplementary Methods

#### Generation and maintenance of Vpr transgenic mice

PEPCK/tTA (T8 line) mice were crossed with tet-op/Vpr mice (L2 line) to generate dual-transgenic mice (Vpr Tg mice). Seven-to-nine week old female WT and Vpr Tg mice were maintained on a diet containing doxycycline (TD 98186, Envigo, Madison, WI) to inhibit the expression of Vpr. Mice were switched from doxycycline diet to 65% protein food (TD 190088, Envigo) for two weeks to induce transgene promoter activity. Both WT and Vpr Tg mice were fed with freshly prepared semi-solid sodium-deficient food [TD 90228 (3 g), containing casein (33.3%), agar (0.5%), sodium (0.045%), and 2.67 mL of water] for four days.(1, 2) Two WT FVB/N mice maintained on regular laboratory chow were also included for comparison. The mouse experimental protocol was approved in advance by the NIDDK Animal Care and Use Committee.

#### Tissue microarray preparation

Seven WT and thirteen Vpr Tg mouse kidney tissues were fixed with 10% buffered formalin for 24 hours, stored in 70% ethanol, and embedded in paraffin. Tissue microarrays (TMAs) containing 5 µm sections of kidney cortex tissues of WT and Vpr Tg mice were prepared at the Experimental Pathology Laboratory, Laboratory of Pathology, Center for Cancer Research, NCI, NIH. Triplicate cortex samples from seven WT and thirteen Vpr Tg mice were placed on the TMA slides. *In situ* hybridization, immunohistochemistry (IHC) and tunnel assay were performed on the TMA slides, as described below.

#### *In situ* hybridization

Fluorescence *in situ* hybridization was performed on TMAs using RNAscope reagents (Advanced Cell Diagnostics, Bio-techne, Minneapolis, MN). Briefly, the TMA slides were de-paraffinized, boiled with RNAscope target retrieval reagent for 15 min and digested with protease at 40 °C for 30 min. This was followed by hybridization for 2 h at 40 °C with RNA probes against *Pvalb* and *Slc12a3*. In addition, *Slc8a1*, *Trpv5* and *Hipk2* were also tested. Specific probe binding sites were visualized using fluorescent RNAscope Hiplex12 Reagents Kit (488, 550, 650) v2 (Catalog # 324419, Advanced Cell Diagnostics).

#### Immunohistochemistry (IHC) of parvalbumin

Tissue microarray slides were deparaffinized and rehydrated; antigen retrieval was performed by treating with citrate-buffered medium for 15 min in a hot water bath. The tissue samples were blocked with 2.5% normal horse serum for 20 min. The sections were incubated for 1 h at room temperature with the primary antibody against DCT1-specific parvalbumin (ab11427, 1 µg/ml, Abcam, Cambridge, UK) which was detected with the ImmPRESS HRP Universal Antibody (horse anti-mouse/rabbit IgG) Polymer Detection Kit and ImmPACT DAB EqV Peroxidase (HRP) Substrate (Vector Laboratories, Burlingame, CA) protocol, and counter stained with hematoxylin.

#### Tunnel assay to detect apoptotic cells

Terminal deoxynucleotidyl transferase dUTP nick end labeling (TUNEL) was used for detecting DNA fragments generated during apoptosis. Tunnel assay on TMA was performed by Histoserve (Germantown, MD).

**Quantification of IHC and TUNEL images**

Stained TMA slides were scanned at 40x on a Hamamatsu Nanozoomer. Using QuPATH (0.3.2), TMA masks were generated following color normalization and tissue detection. For paravalbumin detection, annotation masks for paravalbumin positive and negative tubular profiles were established to allow for automated detection. Percent parvalbumin-positive tubular area ($mm^2$) was calculated as percent of cortical area. For TUNEL quantification, nuclei were segmented and a threshold set for TUNEL positive nuclei, percentage of TUNEL-positive nuclei was calculated.

**Confocal microscopy & image analysis**

A Yokogawa CSU-W1 SoRa spinning disk confocal scan head (Yokogawa, Sugar Land, TX), with 50 micron pinhole (standard confocal mode, no SoRa), mounted on a Nikon Ti2 microscope (Tokyo, Japan). NIS-elements 5.21.02 imaging software (Nikon Instruments, Melville, NY), was used to collect tiles of multi-color fluorescence images. Fluorescence image channels were obtained sequentially, while sharing the Yokogawa T405/488/568/647 dichroic beamspitter.

DAPI fluorescence was excited by the 405nm laser and emission was filtered by ET455/58 (Chroma, Technology Corp, Bellows Falls, VT). Green fluorescence was excited by the 488nm laser and emission was filtered by ET520/40 (Chroma). Orange fluorescence was excited by the 561nm laser and emission was filtered by ET605/52 (Chroma). Far red fluorescence was excited by the 640nm laser and emission was filtered by ET655LP (Chroma). Images used for quantitation of stain prevalence were acquired with the Nikon Plan-Apoλ 20x/0.75 objective lens, producing a confocal section thickness of 7-7.5μm for all fluorescence channels. Whenever images of the same channel were to be compared during analysis, the same settings were used for acquisition.

For pre-processing images, NIS Elements was used to stitch, align, crop, and denoise (AI.denoise) images, followed by export to OME TIF for further processing. Fiji software(3) protocol used to quantify secondary antibody fluorescence occurring in each TMA spot was as follows: 1) Spot ROIs (Regions of Interest) were defined manually to exclude dust and any other potential areas of artifact, and also including an appropriate area around the spot for the auto-thresholds to work properly; 2) Relevant signal area, determined by Triangle threshold, was determined; 3) Nuclear area was determined by Otsu threshold, removing unrealistically small size nuclei from the threshold mask; 4) Signal area calculated above was normalized to tissue area by dividing by nuclear area. Macros were used to facilitate the protocol. For presentation, images were contrast-enhanced, by the same amount when comparing different conditions.

Fiji software was used for generating measurements for each TMA spot, which were copied to a spreadsheet for calculating (Total signal area) / (Total nuclear area).

1. The multi-channel OME TIF images were loaded into Fiji, one TMA at a time. To conserve memory, the BioFormats plugin was used to limit the channels loaded to those

that were currently needed, i.e. the two channels representing the nuclear stain and the signal of interest.

2. A ROI (Region Of Interest) was defined for every spot.
    a. The spot was zoomed to fill the computer monitor, on both channels, to identify and avoid potential artifacts when drawing the ROI. The channel acquired with UV or violet excitation, which is also the channel used to label nuclei in this paper, was particularly useful at revealing contaminant (and autofluorescent) dust.
    b. If the spot was mostly circular and free of artifacts; then a circle was used to define the ROI; otherwise, a polygon, to avoid the artifacts..
    c. ROIs were generally slightly larger than spot boundaries, without forgetting that too many non-biological pixels in the ROI could potentially cause problems later with auto-thresholding.
3. ROI Manager was used to organize the list of ROIs for each TMA, which could be saved for later use, and edited if needed. For convenience, the names of the ROIs were defined according to row and column number, e.g. R1C1, and sorted alphabetically. ROI Manager is required for the two macros described below.

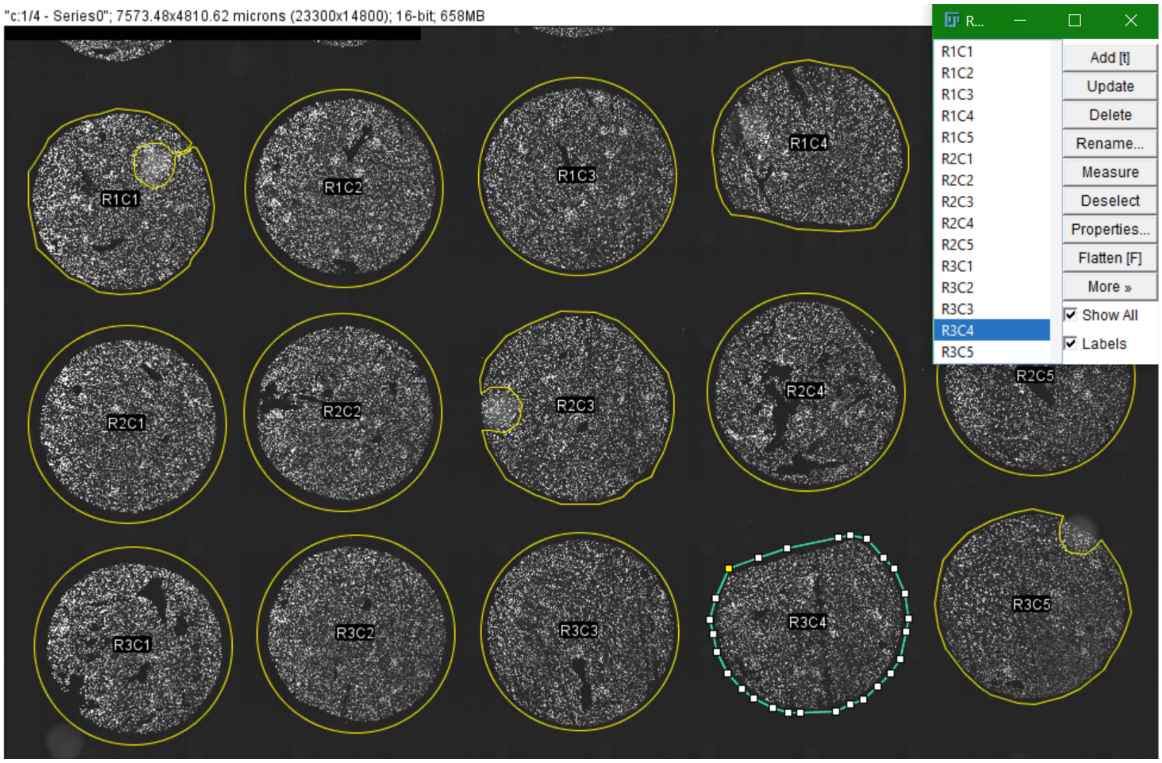


4. Macro "ROIManLoop_Thresh-MeasureSigArea.ijm" measures the total area (in sq. microns) in each TMA spot where valid signal exists.
    a. The Triangle auto-threshold is used to define which pixels qualify as valid signal. The threshold is applied locally to each ROI, to account for deviations in background across the TMA.

```
// Macro ROIManLoop_Thresh-MeasureSigArea.ijm
// Purpose: loops through the ROIs in ROI Manager,
```

```
// ...measuring the area in each ROI that is above threshold (valid signal).
// NOTE: ROIs need to be defined in ROI Manager for this macro to work!!

waitForUser("Please make sure you have the image selected \n for MEASURING
AREA OF FLUORESCENCE SIGNAL, \n and \n ROI Manager has the right
ROI list.\n---> THEN Click OK or press the ENTER key to continue");
currimage=getTitle();
run("Set Measurements...", "area limit display redirect=None decimal=0");
// mean not used for analysis, but could be
Table.create("Results");  // clears Results table
run("Input/Output...", "jpeg=90 gif=-1 file=.csv save_column save_row");
// omitting column and row names for copying data from Results tables

n = roiManager("count");
for (i = 0; i < n; i++) {
   selectWindow(currimage);
   roiManager("select", i);
// Duplication not necessary for measuring, but useful for showing thresholds for
QC,
// plus threshold display is quicker on smaller images.
    RoiImage = Roi.getName+"_SigThr";
    run("Duplicate...", " ");
    setAutoThreshold("Triangle dark");
    // particular auto-threshold method is sample- and probe-dependent
    run("Measure");
    rename(RoiImage);
}
ResultsNewName = "Results for Fluorescence Signal in " + currimage;
Table.rename("Results", ResultsNewName);
run("Tile");
```

b. A cropped image of each spot, with threshold visualized, was generated along with its measurement. These spot-images provided a quick visual QC check, for two things: areas of artifact missed when defining the ROI earlier; and the actual threshold, whether it seems reasonable.
c. If there were any thresholded areas that looked like artifacts, missed during the previous step when defining ROIs, those ROIs could be adjusted.
d. If a spot's auto-threshold seemed to be conspicuously lower that what the viewer's common sense judgement would dictate for "valid signal", the ROI boundaries for those outlier spots are checked to see if too many non-biological background pixels were included. The Triangle threshold uses peaks in the histogram for determining the threshold; if there is measurable autofluorescence above the level of non-biological background, it should be the main source of the histogram peak, not the non-biological background.

   e. If any adjustments were made to the ROIs, the ROI list was re-saved in ROI Manager, and the macro was re-run to generate new Results.
   f. When the visual QC check is passed, the Results are copied to the spreadsheet that has been set up for analysis.

5. Macro “ROIManLoop_Thresh-MeasureNucArea.ijm” measures the total area (in sq. microns) in each TMA spot where nuclear signal exists.
   a. The Otsu auto-threshold was used to define which pixels qualify as nuclear signal. The threshold was applied locally to each ROI, to account for deviations in background across the TMA.

```
// Macro ROIManLoop_Thresh-MeasureNucArea.ijm
// Purpose: loops through the ROIs in ROI Manager,
// ...measuring the area in each ROI that is above threshold (valid nuclear signal).
// Analyze Particles is used to eliminate particles too small to be nuclei.
// NOTE: ROIs need to be defined in ROI Manager for this macro to work!!

waitForUser("Please make sure you have the image selected for MEASURING
NUCLEAR AREA, \n and \n ROI Manager has the right ROI list.\n---> THEN
Click OK or press the ENTER key to continue");
currimage=getTitle();
run("Set Measurements...", "area limit display redirect=None decimal=0");
run("Input/Output...", "jpeg=90 gif=-1 file=.csv save_column save_row");
// omitting column and row names for copying data from Results tables
Table.create("Summary");  // clears Summary table

n = roiManager("count");
for (i = 0; i < n; i++) {
   selectWindow(currimage);
    roiManager("select", i);
// Duplication not necessary for measuring, but useful for creating mask images
for QC
    RoiName = Roi.getName;
    run("Duplicate...", "title=[ROIduplicate]");
    rename(RoiName);
   setAutoThreshold("Otsu dark");
    run("Analyze Particles...", "size=10-Infinity show=Masks exclude clear
include summarize");
    NucMaskImage = RoiName+"_Nucmask";
    rename(NucMaskImage);
    close(RoiName);
}
LatestSummary = "Summary for "+currimage;
Table.rename("Summary", LatestSummary);
run("Tile");
```

b. A cropped image of each spot, converted to an inverted mask of thresholded nuclei, was created as a quick visual QC check.
c. If the nuclear masks (i.e. thresholds) looked reasonable, the Summary table was copied to the spreadsheet for analysis.

***In vitro* detection of apoptosis in Vpr-treated 209 mDCT cells**

Caspases are *c*ysteine-*a*spartic acid-*s*pecific *p*rote*ases* that are activated in response to different cell death-inducing stimuli (1).Therefore, The CHEMICON CaspaTag$^{TM}$ Pan-Caspase *In Situ* Assay Kit (APT420, EMD Millipore) was used according to the manufacturer's protocol, with minor changes as described below to detect apoptosis induced in mDCT cells treated with sVpr.

The 209 mDCT cells (ATCC) were grown in DMEM containing 10% FBS and penicillin-streptomycin. For the assay, 10,000 cells /200 µl of culture medium were grown overnight in a 96-well plate. Next day, cells were treated with opti medium containing sVpr (1ng/200µl/well). The control well received Opti-MEM medium (Gibco) only. Twenty four hours later, the cells were stained with 1x FLICA for 1hour at 370C, then washed twice with 1x wash buffer provided with the kit. Cells were trypsinized with 50µl of 0.025% trypsin and neutrilized with 50µl of medium. Accutase (Cat.# 00-4555-56 Thermo Fisher) containing propidium iodide *(*PI 250ng/100µl accutase) was added, immediately before the FACS analysis. Cells were then viewed at excitation at 490nm, emission >520nm. PI has a maximum emission of 637nm.

**Quantitive PCR of *Ier3* in 209 mDCT cells**

Mouse 209 DCT cells ($10^5$ /ml ) were grown overnight in six well plates. Next day, cells were treated with fresh medium containing 0.0. 0.1,10 and 100 ng/ml of sVpr for 24 hours. The RNA from control and Vpr-treated 209 mDCT cells were prepared by Tryzol (Sigma). The forward and reverse primers used for Ier3 amplification were: mmu-Ier3F ACACCTGAGCCCATTTCTG and mmu-Ier3-R TGACCCATCGCGTTTAGAAG, respectively. The values were normalized to beta-actin amplified using the following primers: mActb-F CCACCATGTACCCAGGCATT and mActb-R AGGGTGTAAAACGCAGCTCA.

### Supplementary Figures

(a)

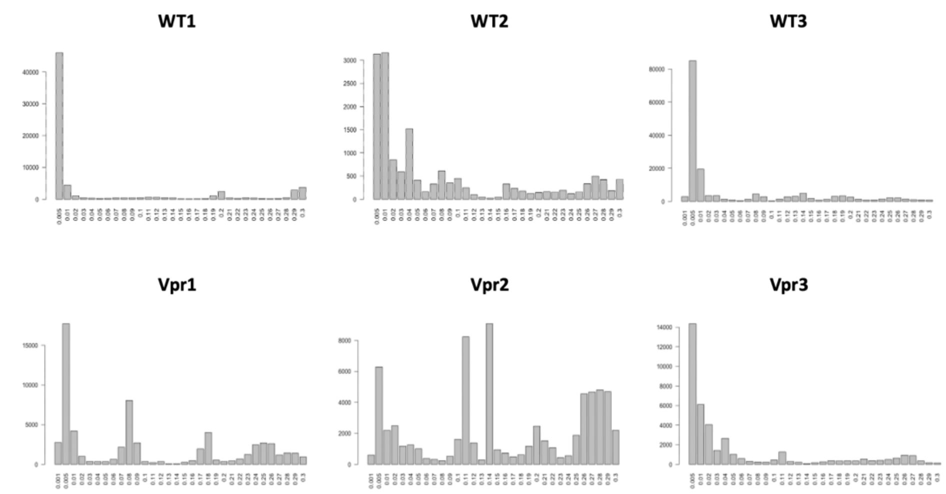


(b)

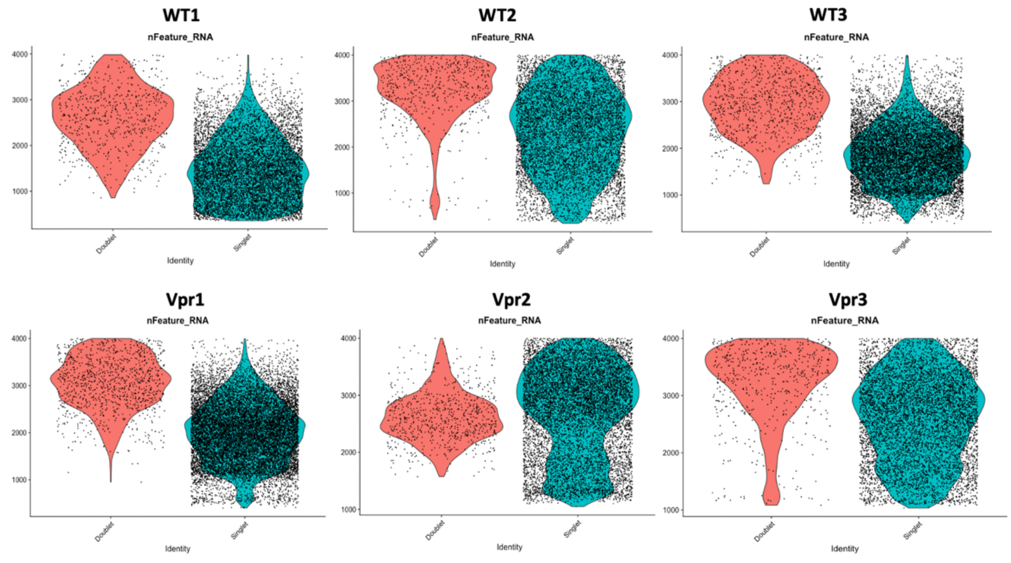


**Supplementary Figure 1.** Results of doublet detection analysis in each sample by DoubletFinder. (a) Barplots showing BC metrices (Y-axis) for different pK values (X-axis). (b) Violin plots showing the nFeature_RNA counts in predicted single and doublet samples.

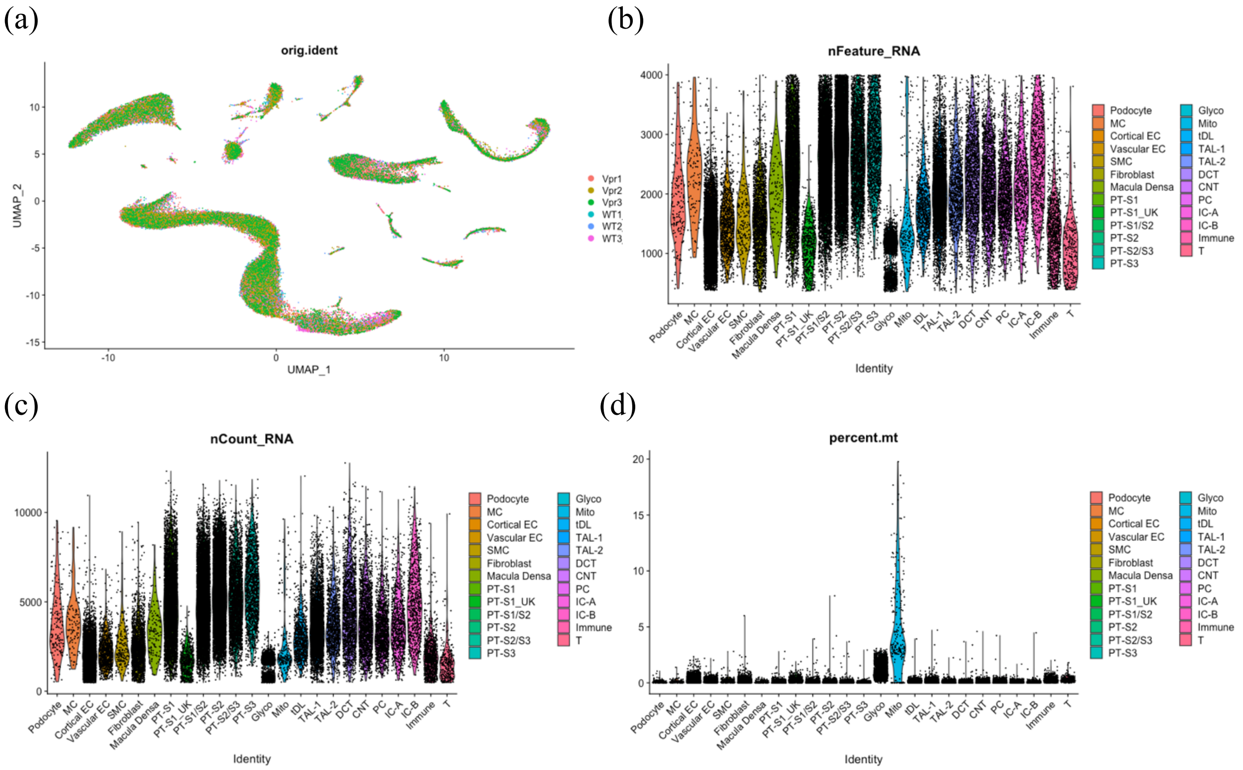


**Supplementary Figure 2.** snRNA-seq analysis of the dataset after anchor-based integration and batch correction by harmony. (a) UMAP of the merged dataset with original samples in different colors (b) Violin plot showing the number of features (genes) expressed in each cell in each cluster (c) Violin plot showing the total number of mRNA expressed in each cell in each cluster (d) Violin plot showing the percentage of mitochondrial transcripts in each cell in each cluster.

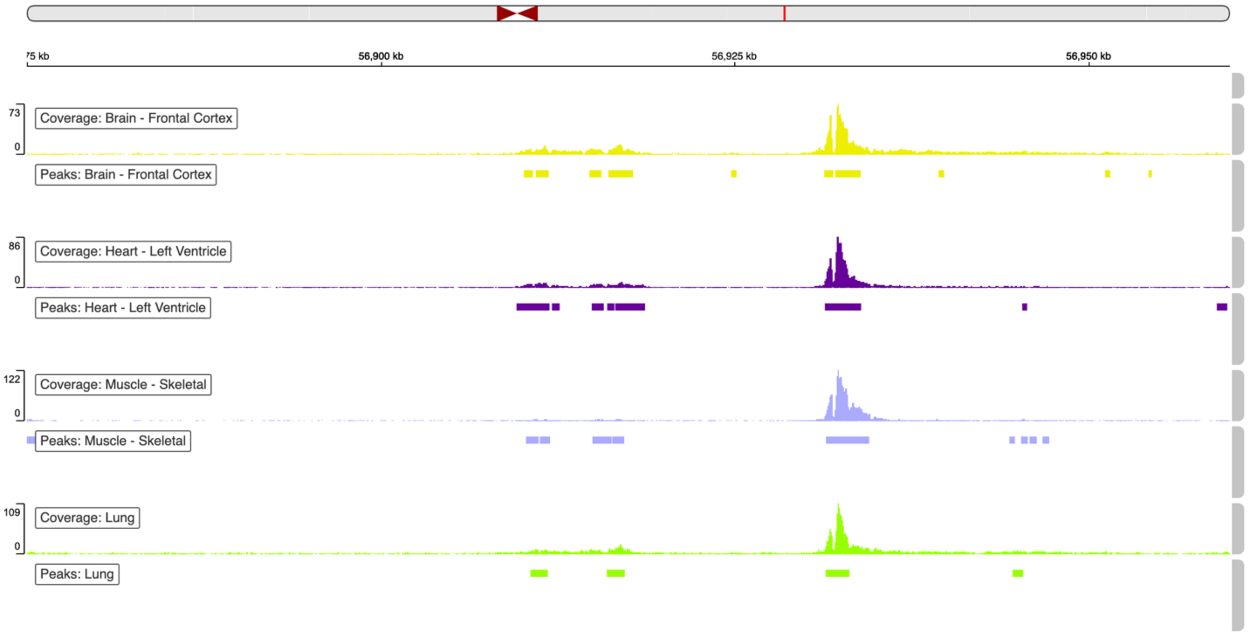


**Supplementary Figure 3.** H3K27ac enhancer mark on human chromosome 16 covering *SLC12A3* region in frontal cortex, left ventricle, skeletal muscle and lung retrieved from GTEx database.

(a)

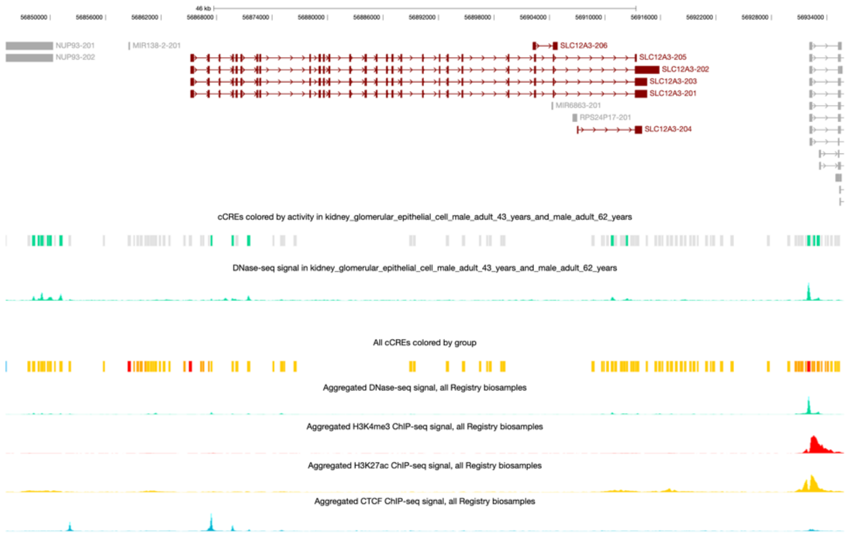


(b)

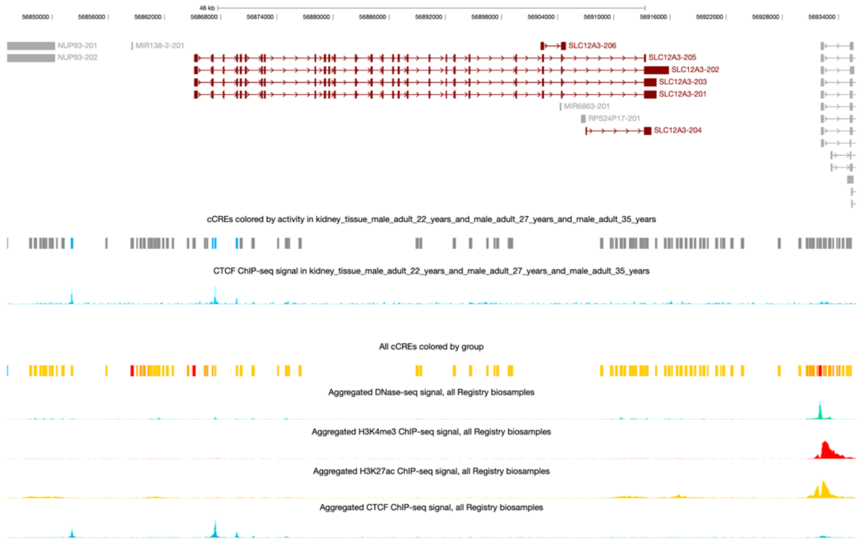


**Supplementary Figure 4.** Candidate cis-regulatory elements (cCREs) in human SLC12A3 region retrieved from ENCODE database showing multiple enhancer-like elements (yellow vertical bars) in all registry samples. (a) DNase-seq signal in glomerular epithelial cells from two adult males (b) CTCF ChIP-seq signal in kidney tissues from three adult males.

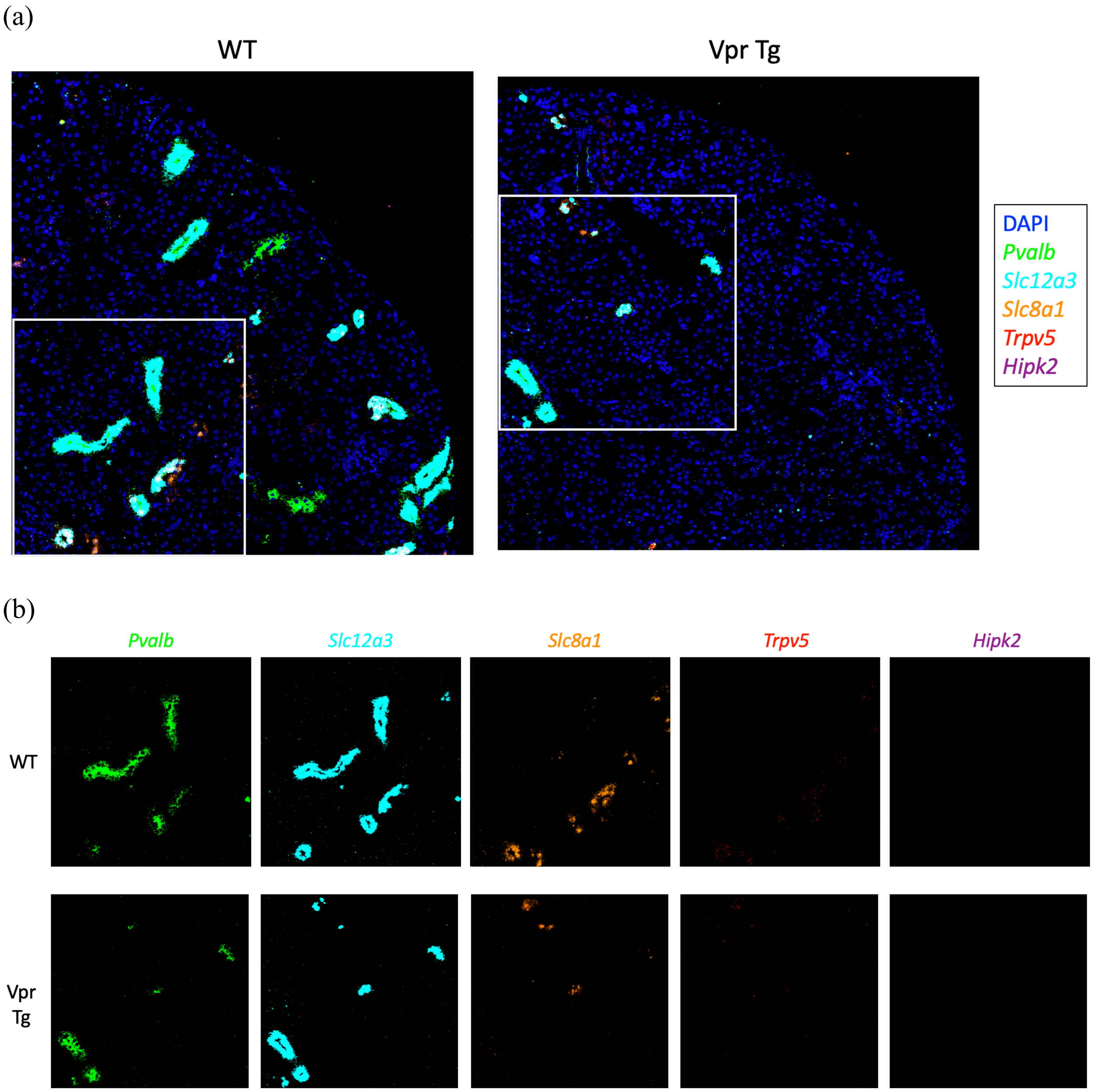


**Supplementary Figure 5.** Imaging results of WT and Vpr Tg cortex samples. (a) RNAScope images of the right upper quadrant of a tissue microarray section showed that compared to WT, Vpr Tg sample had less *Slc12a3* and *Pvalb* fluorescence signals. (b) Areas of images from figure (a) marked by the white rectangles shows individual genes for *Pvalb*, *Slc12a3*, *Slc8a1*, *Trpv5* and *Hipk2*.

(a)

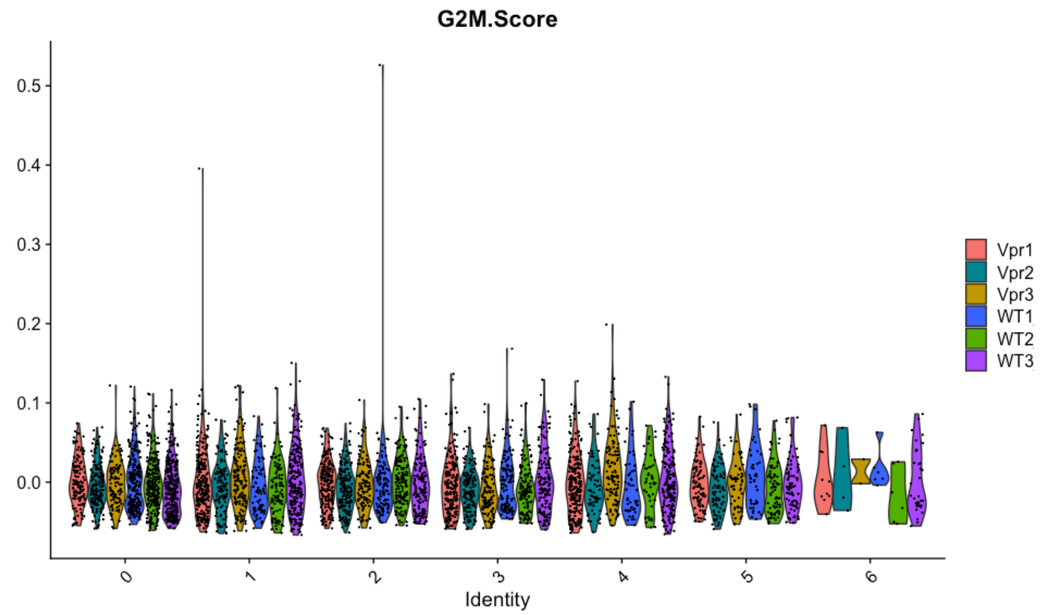


(c)

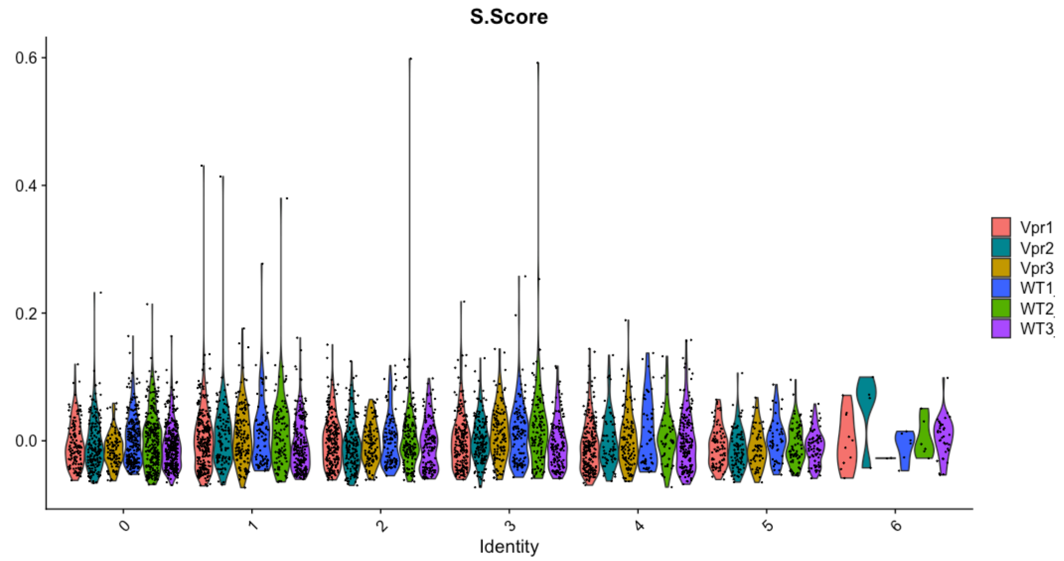


**Supplementary Figure 6.** Violin plots showing the aggregate gene scores of cell cycle genes in distal tubular cell subclusters. (a) G2M scores of 48 G2M phase genes (b) S scores of 40 S phase genes. Subcluster 0 = DCT1; subcluster 5 = DCT2; subcluster 6 = *Nr3c2*$^{+}$ DCT.

| **Cluster** | **WT1** | **WT2** | **WT3** | **Vpr1** | **Vpr2** | **Vpr3** |
|---|---|---|---|---|---|---|
| Podocyte | 0.39 | 0.44 | 0.37 | 0.44 | 0.05 | 0.13 |
| MC | 0.11 | 0.37 | 0.19 | 0.17 | 0.30 | 0.01 |
| Cortical EC | 13.68 | 14.11 | 9.49 | 10.78 | 12.86 | 12.65 |
| Vascular EC | 1.49 | 1.51 | 1.85 | 1.83 | 1.56 | 2.34 |
| Fibroblast | 4.22 | 3.53 | 2.96 | 3.11 | 4.89 | 2.80 |
| SMC | 0.26 | 0.33 | 0.44 | 0.50 | 0.20 | 0.40 |
| Macula Densa | 0.23 | 0.27 | 0.19 | 0.27 | 0.22 | 0.37 |
| PT-S1 | 10.80 | 10.26 | 9.96 | 8.93 | 13.06 | 8.22 |
| PT-S1/S2 | 14.37 | 14.57 | 14.99 | 14.49 | 21.96 | 13.34 |
| PT-S2 | 22.34 | 20.05 | 11.00 | 11.52 | 16.13 | 14.12 |
| PT-S2/S3 | 5.93 | 5.43 | 5.27 | 6.34 | 2.84 | 6.37 |
| PT-S3 | 3.43 | 2.02 | 3.91 | 2.40 | 1.75 | 2.56 |
| PT-S1_UK | 0.95 | 1.25 | 0.73 | 1.30 | 0.55 | 1.49 |
| Glyco | 1.29 | 2.61 | 4.89 | 5.62 | 6.69 | 2.86 |
| tDL | 0.39 | 0.33 | 3.00 | 2.41 | 0.16 | 2.06 |
| TAL-1 | 4.03 | 4.46 | 13.69 | 13.20 | 2.69 | 13.38 |
| TAL-2 | 1.39 | 1.71 | 2.54 | 2.72 | 1.15 | 2.74 |
| **DCT** | **3.58** | **4.30** | **3.67** | **1.86** | **2.86** | **2.36** |
| CNT | 2.94 | 3.84 | 3.00 | 3.53 | 3.57 | 3.77 |
| PC | 1.76 | 2.11 | 3.33 | 2.87 | 2.03 | 3.29 |
| IC-A | 1.16 | 1.31 | 1.92 | 1.38 | 0.51 | 1.77 |
| IC-B | 0.88 | 1.43 | 1.16 | 1.26 | 1.43 | 1.49 |
| Immune | 2.44 | 2.55 | 0.99 | 2.51 | 1.24 | 1.09 |
| T | 1.18 | 0.68 | 0.22 | 0.50 | 0.39 | 0.21 |
| Mito | 0.76 | 0.53 | 0.25 | 0.05 | 0.92 | 0.18 |
| **DCT subclusters** | | | | | | |
| **DCT1** | **3.03** | **3.41** | **2.87** | **1** | **1.93** | **1.47** |
| DCT2 | 0.58 | 1.02 | 0.68 | 0.64 | 0.95 | 0.83 |
| *Nr3c1*$^{+}$ DCT | 0.08 | 0.08 | 0.31 | 0.07 | 0.04 | 0.03 |

**Supplementary Table 1**. Metrics of the 25 cells clusters and 3 DCT sub-clusters that were identified in the unsupervised clustering of six mouse renal cortex samples. The percentage of individual cell type cells in each sample was calculated from the total number of cells from that sample. MC mesangial cell; Cortical EC = cortical endothelial cell; Vascular EC = vascular endothelial cell; SMC = smooth muscle cell; PT = proximal tubule; Glyco = Cells with high expression of glycolytic enzymes; tDL = thin descending segment; TAL = thick ascending limb; DCT = distal convoluted tubule; CNT = connecting tubule; PC = principal cell; IC = intercalated cell; T = T cell; Mito = mitochondria-enriched cell.

| Variant Id | SNP Id | P-Value | NES | Tissue |
|---|---|---|---|---|
| chr16_56715244_C_T_b38 | rs11649088 | 3.80E-05 | -0.27 | Spleen |
| chr16_56717614_G_T_b38 | rs9932268 | 3.90E-06 | -0.45 | Spleen |
| chr16_56718282_C_CA_b38 | rs34867225 | 9.80E-07 | 0.33 | Spleen |
| chr16_56719719_C_T_b38 | rs12931136 | 1.10E-06 | 0.32 | Spleen |
| chr16_56721606_A_G_b38 | rs12923723 | 2.80E-06 | 0.32 | Spleen |
| chr16_56723904_A_G_b38 | rs4783955 | 2.50E-05 | 0.27 | Spleen |
| chr16_56729100_A_T_b38 | rs9939873 | 2.50E-05 | -0.28 | Spleen |
| chr16_56730563_A_G_b38 | rs4784726 | 4.20E-06 | 0.35 | Spleen |
| chr16_56732919_C_T_b38 | rs4783956 | 2.60E-05 | 0.28 | Spleen |
| chr16_56733179_G_A_b38 | rs12934065 | 1.10E-06 | 0.32 | Spleen |
| chr16_56733213_T_A_b38 | rs12921489 | 1.10E-06 | 0.32 | Spleen |
| chr16_56733890_C_T_b38 | rs12921698 | 1.10E-06 | 0.32 | Spleen |
| chr16_56737879_T_C_b38 | rs62036970 | 3.10E-05 | 0.28 | Spleen |
| chr16_56746479_T_C_b38 | rs12930689 | 2.60E-05 | 0.28 | Spleen |
| chr16_56751211_C_A_b38 | rs73555563 | 3.90E-06 | -0.45 | Spleen |
| chr16_56754842_G_A_b38 | rs12930414 | 1.10E-06 | 0.32 | Spleen |
| chr16_56758087_T_G_b38 | rs12924698 | 3.40E-05 | 0.25 | Spleen |
| chr16_56766421_T_A_b38 | rs8048885 | 1.10E-06 | -0.32 | Spleen |
| chr16_56768702_TA_T_b38 | rs564962218 | 1.90E-07 | -0.73 | Spleen |
| chr16_56774700_T_C_b38 | rs4371151 | 2.80E-05 | -0.27 | Spleen |
| chr16_56775541_AG_A_b38 | rs60454311 | 4.00E-05 | -0.27 | Spleen |
| chr16_56777549_A_C_b38 | rs9928890 | 3.90E-06 | -0.45 | Spleen |
| chr16_56780935_G_A_b38 | rs8052978 | 1.50E-05 | -0.26 | Spleen |
| chr16_56784635_T_G_b38 | rs6499852 | 6.60E-07 | -0.33 | Spleen |
| chr16_56785259_T_G_b38 | rs6416772 | 6.60E-07 | -0.33 | Spleen |
| chr16_56786728_G_A_b38 | rs7500207 | 1.60E-05 | -0.28 | Spleen |
| chr16_56792854_T_C_b38 | rs9938953 | 6.40E-07 | -0.3 | Spleen |
| chr16_56794811_C_T_b38 | rs13335111 | 4.00E-05 | -0.27 | Spleen |
| chr16_56795628_G_A_b38 | rs12932037 | 1.50E-05 | 0.29 | Spleen |
| chr16_56796867_G_C_b38 | rs10048067 | 1.50E-05 | 0.29 | Spleen |
| chr16_56797234_C_T_b38 | rs9922951 | 4.00E-05 | -0.27 | Spleen |
| chr16_56798390_C_T_b38 | rs9927884 | 3.90E-06 | -0.45 | Spleen |
| chr16_56800184_A_G_b38 | rs4783959 | 2.60E-06 | -0.28 | Spleen |
| chr16_56801800_G_T_b38 | rs8051691 | 4.00E-05 | -0.27 | Spleen |
| chr16_56802196_T_C_b38 | rs9929577 | 5.20E-06 | -0.27 | Spleen |
| chr16_56804675_G_A_b38 | rs7189328 | 1.60E-05 | 0.28 | Spleen |
| chr16_56805066_T_G_b38 | rs13338063 | 4.70E-05 | -0.25 | Spleen |
| chr16_56806259_A_G_b38 | rs7187512 | 8.40E-07 | -0.29 | Spleen |

| | | | | |
|---|---|---|---|---|
| chr16_56806416_G_A_b38 | rs10221121 | 1.30E-05 | 0.29 | Spleen |
| chr16_56807802_C_T_b38 | rs9938980 | 2.60E-05 | -0.25 | Spleen |
| chr16_56810727_TACCAGCGCCCCAGAAATCCTCTTTGTGGTCCTTTCTGGTC_T_b38 | rs141700288 | 1.70E-06 | 0.33 | Spleen |
| chr16_56811388_A_T_b38 | rs72786721 | 1.90E-07 | -0.73 | Spleen |
| chr16_56812624_T_C_b38 | rs12925122 | 6.60E-07 | 0.33 | Spleen |
| chr16_56813358_G_A_b38 | rs1529928 | 1.70E-06 | 0.33 | Spleen |
| chr16_56815584_G_A_b38 | rs1529929 | 8.40E-07 | -0.29 | Spleen |
| chr16_56816611_A_G_b38 | rs13338782 | 3.90E-06 | -0.45 | Spleen |
| chr16_56816690_A_G_b38 | rs9939678 | 8.40E-07 | -0.29 | Spleen |
| chr16_56818477_T_C_b38 | rs12928581 | 1.60E-05 | 0.28 | Spleen |
| chr16_56818910_G_T_b38 | rs1561139 | 1.20E-06 | -0.29 | Spleen |
| chr16_56820372_C_T_b38 | rs12930486 | 1.60E-05 | 0.28 | Spleen |
| chr16_56825304_C_T_b38 | rs12919839 | 1.50E-05 | 0.29 | Spleen |
| chr16_56825683_G_A_b38 | rs7199480 | 8.40E-07 | -0.29 | Spleen |
| chr16_56828083_C_G_b38 | rs2099536 | 3.70E-07 | -0.3 | Spleen |
| chr16_56828231_GA_G_b38 | rs34735016 | 3.70E-07 | -0.3 | Spleen |
| chr16_56830486_C_T_b38 | rs1561140 | 5.20E-05 | -0.25 | Spleen |
| chr16_56830706_C_T_b38 | rs4461062 | 8.40E-06 | -0.27 | Spleen |
| chr16_56831337_C_A_b38 | rs4784730 | 8.40E-06 | -0.27 | Spleen |
| chr16_56832590_T_C_b38 | rs12918918 | 6.60E-07 | 0.33 | Spleen |
| chr16_56832889_G_T_b38 | rs6499853 | 1.10E-06 | 0.32 | Spleen |
| chr16_56834788_A_G_b38 | rs1347591 | 1.20E-05 | -0.27 | Spleen |
| chr16_56836171_G_GT_b38 | rs35892526 | 3.00E-06 | 0.31 | Spleen |
| chr16_56837557_TTTTATTGATTAC_T_b38 | rs145442551 | 3.90E-06 | -0.45 | Spleen |
| chr16_56838810_C_T_b38 | rs3764265 | 3.90E-06 | -0.45 | Spleen |
| chr16_56840285_G_A_b38 | rs1865830 | 8.40E-07 | -0.29 | Spleen |
| chr16_56840945_C_T_b38 | rs2007432 | 8.40E-07 | -0.29 | Spleen |
| chr16_56841360_A_AT_b38 | rs3214653 | 8.40E-07 | -0.29 | Spleen |
| chr16_56845290_T_C_b38 | rs12918087 | 9.80E-06 | 0.33 | Spleen |
| chr16_56845659_G_C_b38 | rs7205421 | 3.60E-07 | 0.3 | Spleen |
| chr16_56845726_G_C_b38 | rs35172527 | 3.60E-06 | 0.31 | Spleen |
| chr16_56846187_GATCACC_G_b38 | rs147271366 | 1.40E-06 | 0.31 | Spleen |
| chr16_56846246_A_G_b38 | rs2399562 | 3.90E-07 | -0.3 | Spleen |
| chr16_56846493_T_C_b38 | rs12929119 | 1.40E-06 | 0.31 | Spleen |
| chr16_56846969_AGGTTCTTTGACCTGTAGATG_A_b38 | rs144725437 | 1.40E-06 | 0.31 | Spleen |
| chr16_56848952_AAG_A_b38 | rs72067847 | 9.80E-06 | 0.33 | Spleen |
| chr16_56849526_G_A_b38 | rs8044753 | 5.50E-05 | -0.25 | Spleen |
| chr16_56849797_G_C_b38 | rs8045306 | 8.40E-07 | -0.29 | Spleen |

| | | | | |
|---|---|---|---|---|
| chr16_56850012_A_G_b38 | rs735144 | 8.40E-07 | -0.29 | Spleen |
| chr16_56851226_T_G_b38 | rs6499855 | 7.10E-06 | -0.27 | Spleen |
| chr16_56854080_G_A_b38 | rs76942671 | 1.90E-07 | -0.73 | Spleen |
| chr16_56856531_C_CA_b38 | rs35810185 | 6.00E-05 | 0.3 | Spleen |
| chr16_56860192_T_C_b38 | rs12933363 | 4.00E-06 | 0.3 | Spleen |
| chr16_56860593_G_T_b38 | rs12932041 | 3.00E-06 | 0.31 | Spleen |
| chr16_56861122_G_T_b38 | rs1436424 | 6.30E-07 | -0.3 | Spleen |
| chr16_56862124_T_C_b38 | rs12599065 | 7.80E-07 | -0.3 | Spleen |
| chr16_56862277_A_G_b38 | rs12921781 | 2.30E-06 | 0.31 | Spleen |
| chr16_56862818_G_A_b38 | rs3829502 | 9.70E-08 | -0.32 | Spleen |
| chr16_56866607_C_T_b38 | rs12918664 | 3.20E-06 | 0.29 | Spleen |
| chr16_56871338_C_T_b38 | rs1123507 | 2.10E-13 | -0.68 | Spleen |
| chr16_56871338_C_T_b38 | rs1123507 | 2.50E-06 | -0.33 | Small Intestine - Terminal Ileum |
| chr16_56873040_CT_C_b38 | rs55822178 | 2.20E-12 | -0.64 | Spleen |
| chr16_56873040_CT_C_b38 | rs55822178 | 2.50E-05 | -0.31 | Small Intestine - Terminal Ileum |
| chr16_56893416_A_C_b38 | rs28728226 | 3.40E-05 | 0.32 | Pancreas |
| chr16_56893529_G_C_b38 | rs16963397 | 3.40E-05 | 0.32 | Pancreas |
| chr16_56893808_T_C_b38 | rs12449167 | 3.40E-05 | 0.32 | Pancreas |
| chr16_56893846_C_A_b38 | rs12445576 | 3.30E-05 | 0.32 | Pancreas |
| chr16_56894038_C_T_b38 | rs12445625 | 3.30E-05 | 0.32 | Pancreas |
| chr16_56894148_T_C_b38 | rs12449249 | 3.40E-05 | 0.32 | Pancreas |
| chr16_56894165_A_G_b38 | rs12448598 | 3.30E-05 | 0.32 | Pancreas |
| chr16_56894270_G_A_b38 | rs12445505 | 3.30E-05 | 0.32 | Pancreas |
| chr16_56894304_C_T_b38 | rs12445698 | 3.30E-05 | 0.32 | Pancreas |
| chr16_56894368_G_C_b38 | rs13306679 | 3.30E-05 | 0.32 | Pancreas |
| chr16_56894763_C_A_b38 | rs2289118 | 3.30E-05 | 0.32 | Pancreas |
| chr16_56894793_A_C_b38 | rs9921308 | 3.30E-05 | 0.32 | Pancreas |
| chr16_56895768_G_T_b38 | rs13338222 | 3.00E-05 | 0.32 | Pancreas |
| chr16_56895929_T_C_b38 | rs13334864 | 3.70E-05 | 0.32 | Pancreas |
| chr16_56895989_A_G_b38 | rs9929395 | 3.10E-05 | 0.32 | Pancreas |
| chr16_56896007_C_T_b38 | rs9939276 | 3.00E-05 | 0.32 | Pancreas |
| chr16_56896008_T_G_b38 | rs9931565 | 3.10E-05 | 0.32 | Pancreas |
| chr16_56896317_G_T_b38 | rs8056954 | 3.00E-05 | 0.32 | Pancreas |
| chr16_56896339_T_C_b38 | rs8063291 | 2.20E-05 | 0.33 | Pancreas |
| chr16_56896519_A_G_b38 | rs8063278 | 3.00E-05 | 0.32 | Pancreas |
| chr16_56896638_G_A_b38 | rs8061631 | 5.80E-06 | 0.34 | Pancreas |

| | | | | |
|---|---|---|---|---|
| chr16_56896723_G_A_b38 | rs8061810 | 3.00E-05 | 0.32 | Pancreas |
| chr16_56896753_T_A_b38 | rs8047432 | 3.00E-05 | 0.32 | Pancreas |
| chr16_56897008_T_C_b38 | rs34589259 | 2.70E-05 | 0.32 | Pancreas |
| chr16_56897136_G_C_b38 | rs16963412 | 2.70E-05 | 0.32 | Pancreas |
| chr16_56897409_A_G_b38 | rs12923922 | 2.70E-05 | 0.32 | Pancreas |
| chr16_56897553_T_A_b38 | rs12447990 | 5.80E-06 | 0.34 | Pancreas |
| chr16_56897653_C_T_b38 | rs7188963 | 5.80E-06 | 0.34 | Pancreas |
| chr16_56897792_G_A_b38 | rs7187932 | 1.10E-05 | 0.33 | Pancreas |
| chr16_56898410_T_C_b38 | rs13338836 | 1.10E-05 | 0.33 | Pancreas |
| chr16_56898619_G_A_b38 | rs34433002 | 1.10E-05 | 0.33 | Pancreas |
| chr16_56899308_A_G_b38 | rs12448372 | 1.10E-05 | 0.33 | Pancreas |
| chr16_56901178_G_C_b38 | rs6499857 | 3.40E-05 | 0.31 | Pancreas |
| chr16_56904610_C_T_b38 | rs2289114 | 8.90E-06 | 0.34 | Pancreas |
| chr16_56933265_T_G_b38 | rs72786778 | 5.50E-07 | -0.83 | Spleen |
| chr16_56933265_T_G_b38 | rs72786778 | 9.10E-06 | -0.43 | Small Intestine - Terminal Ileum |
| chr16_57405846_A_T_b38 | rs62037105 | 3.40E-05 | -0.48 | Lung |
| chr16_57407454_T_A_b38 | rs60679405 | 3.40E-05 | -0.48 | Lung |
| chr16_57408777_T_C_b38 | rs11218679<br>4 | 3.40E-05 | -0.48 | Lung |
| chr16_57409259_G_A_b38 | rs62037108 | 3.10E-05 | -0.48 | Lung |
| chr16_57410859_A_G_b38 | rs62037110 | 1.80E-05 | -0.53 | Lung |
| chr16_57414449_T_C_b38 | rs59247888 | 3.40E-05 | -0.48 | Lung |
| chr16_57415695_CG_C_b38 | rs14936765<br>9 | 3.40E-05 | -0.48 | Lung |

**Supplementary Table 2**. Expression quantitative trait locus (eQTL) SNPs associated with *SLC12A3* expression levels across different tissues retrieved from the Genotype Tissue Expression (GTEx) database.

Anln
Anp32e
Aurka
Aurkb
Birc5
Bub1
Ccnb2
Cdc20
Cdc25c
Cdca2
Cdca3
Cdca8
Cenpa
Cenpe
Cenpf
Ckap2
Ckap2l
Ckap5
Cks1b
Cks2
Ctcf
Dlgap5
Ect2
G2e3
Gtse1
Hjurp
Hmgb2
Hmmr
Kif11
Kif20b
Kif23
Kif2c
Lbr
Mki67
Ncapd2
Ndc80
Nek2
Nuf2
Nusap1
Psrc1

Rangap1
Smc4
Tacc3
Top2a
Tpx2
Ttk
Tubb4b
Ube2c

**Supplementary Table 3.** The mouse G2M phase genes used to calculate aggregate gene scores shown in **Supplementary Figure 5a**.

Blm
Brip1
Casp8ap2
Ccne2
Cdc45
Cdc6
Cdca7
Chaf1b
Clspn
Dscc1
Dtl
E2f8
Exo1
Fen1
Gins2
Gmnn
Hells
Mcm2
Mcm4
Mcm5
Mcm6
Msh2
Nasp
Pcna
Pola1
Prim1
Rad51
Rad51ap1
Rfc2
Rpa2
Rrm1
Rrm2
Slbp
Tipin
Tyms
Ubr7
Uhrf1
Ung
Usp1
Wdr76

**Supplementary Table 4.** The mouse S phase genes used to calculate aggregate gene scores shown in **Supplementary Figure 5b.**
